# The MUC1 extracellular domain and cytoplasmic tail play distinct roles during *Salmonella* invasion of enterocytes

**DOI:** 10.1101/2025.10.15.682509

**Authors:** Jinyi Su, Xinyue Li, Evelien Floor, Liane ZX Huang, Jos P.M. van Putten, Karin Strijbis

**Affiliations:** Division of Infectious Diseases and Immunology, Department of Biomolecular Health Sciences, Faculty of Veterinary Medicine, Utrecht University, Utrecht, Netherlands

**Keywords:** *Salmonella enterica* serovar Enteritidis, transmembrane mucin, enterocyte invasion, NFκB activation

## Abstract

The intestinal mucus layer consists of secreted and transmembrane (TM) mucins expressed on the apical surface of enterocytes. The TM mucin MUC1 has a highly O-glycosylated extracellular domain (ED) and a cytoplasmic tail (CT) with signaling potential. MUC1 is a target for the *Salmonella* adhesin SiiE, which mediates apical invasion of the bacterium into enterocytes. Here, we determined the contributions of the MUC1 ED and CT to *Salmonella* invasion and subsequent host immune responses. Enzymatic removal of the MUC1 ED from HT29-MTX intestinal cultures blocked *Salmonella* invasion to levels comparable to MUC1 knockout cells. CRISPR-mediated targeted deletion of the MUC1 CT (MUC1-ΔCT) did not quantitatively affect *Salmonella* invasion. To investigate downstream host responses, RNAseq transcriptomics analysis of uninfected and *Salmonella*-infected MUC1-WT, MUC1-ΔCT, and ΔMUC1 cultures was performed. Deletion of full-length MUC1 greatly altered the transcriptome, while only a small group of 132 genes was differentially expressed in MUC1-ΔCT cultures during infection. Several of these CT-dependent genes are related to the NFκB pathway. Immunoblot analysis demonstrates that under uninfected conditions, expression of NFκB subunits RelB, NfkB1-p105, NfkB2-p100, and IκBα was significantly lower in MUC1-WT compared to MUC1-ΔCT and ΔMUC1 cultures. Secretion of cytokines and immune factors was severely reduced in ΔMUC1 cultures, coinciding with reduced *Salmonella* invasion. In MUC1-ΔCT cultures, only galectin-3 and IL-18 secretion were significantly reduced. We conclude that the MUC1 ED is essential for *Salmonella* invasion, while the CT modulates the canonical and non-canonical NFκB pathway, pointing at distinct roles for MUC1 domains in microbe-host interactions and signaling.

**Importance:** The intestinal mucus layer plays an important role in separating commensal and pathogenic microbes from the underlying epithelium. The transmembrane mucin MUC1 is expressed by different types of intestinal epithelial cells and is thought to have important protective and signaling functions. However, enteropathogenic *Salmonella* bacteria can hijack MUC1 through engagement with the SiiE adhesin which leads to bacterial invasion of enterocytes at the apical surface. In this study, we determined how the different MUC1 domains contributed to *Salmonella* invasion and subsequent host responses. We found that the glycosylated MUC1 extracellular domain, but not the cytoplasmic tail, is essential for bacterial invasion. In infected and uninfected intestinal cultures, the MUC1 cytoplasmic tail modulates immune responses including NFκB activation and cytokine secretion. Our study contributes to our understanding of the diverse functions of transmembrane mucins at the intestinal microbe-host interface.

## Introduction

The mucosal surface of the gastrointestinal (GI) tract is covered by a mucus layer that prevents direct interaction of the microbiota with the epithelium and the immune system (1). The mucus layer is composed of a meshwork of mucins that form the main component of the mucosal barrier. There are two main types of mucins, secreted gel-forming mucins and transmembrane mucins that form the glycocalyx (2, 3). Secreted mucins are expressed by goblet cells and released into the gut lumen. Intestinal transmembrane mucins are tethered to the apical surface of epithelial cells and are the first point of contact after pathogens have penetrated the soluble mucus layer (2, 3). MUC1 is one of the most highly expressed transmembrane mucins in the GI tract and the interaction between MUC1 and different microbes has been widely demonstrated (3–5). MUC1 is a type I transmembrane glycoprotein that consists of a very large extracellular domain (ED), a sea urchin sperm protein, enterokinase, and agrin (SEA) module, a transmembrane domain, and a cytoplasmic tail (CT) with signaling potential (6). The MUC1 ED is highly O-glycosylated on the variable number tandem repeat (VNTR) region and extends up to 200 nm from the apical cell membrane into the lumen (7). The SEA domain contains a conserved cleavable GSVV sequence, which enables autoproteolytic cleavage and potential release at the cell surface (8, 9). When present at the epithelial glycocalyx, commensal and pathogenic bacteria can interact with the MUC1 ED. The MUC1 CT consists of 72 amino acids and 22 potential phosphorylation sites (10, 11) and signaling by the CT has been implicated in bacterial infections but also in carcinogenesis and tumor metastasis.

Experiments with the gastric pathogen *Helicobacter pylori* (*H. pylori*) pointed to a conventional role for MUC1 as a host defense molecule in both mouse and human cell models. The large MUC1 ED that protrudes into the lumen likely provides a physical barrier that prevents bacterial contact with the underlying epithelium. Mice lacking Muc1 display a 5-fold higher *H. pylori* colonization and more severe gastritis than wild-type mice (12). Furthermore, co-culture of *H. pylori* with MUC1-expressing gastric epithelial cells induces shedding of MUC1, and shed MUC1 binds to *H. pylori* as a decoy receptor, limiting colonization of the cell surface (13). The role of MUC1 CT in the protection against infection is less well understood. Several studies point towards a general anti-inflammatory effect of MUC1 during activation of TLR5 (14), the NLRP3 inflammasome (15) and during NFκB signaling (16). The NFκB pathway is one of the major pro-inflammatory pathways in the intestinal epithelium, and nuclear translocation of its transcriptional activators is tightly regulated to prevent unnecessary activation. In the canonical NFκB pathway, NFκB1, inhibitory к kinases IKKα, IKKβ, IKKγ and IκBα form different complexes that regulate the inhibition and activation of the transcriptional unit RelA. In the noncanonical NFκB pathway, NFκB2 and IKKα regulate the inhibition and activation of the transcriptional unit RelB. Lipopolysaccharides from the *Salmonella* outer membrane stimulate activation of both NFκB pathways in a temporal manner, where the canonical NFκB pathway is first activated then followed by the noncanonical pathway (17). MUC1 CT was found to interact with IKKγ of the canonical NFκB pathway during infection with *H. pylori*, leading to reduced activation of NF-кB and lower levels of interleukin (IL)-8 (18).

Pathogens can also utilize MUC1 for adhesion or invasion. In respiratory cells, MUC1 serves as a receptor for *Pseudomonas aeruginosa* or its flagellin (19). Direct binding of this pathogen or its flagellin to MUC1 induces phosphorylation of the MUC1 cytoplasmic tail and activation of the mitogen-activated protein (MAP) kinase pathway (20). In intestinal HT29-MTX cultures, *Salmonella* can utilize MUC1 for apical invasion of enterocytes (21). We demonstrated that the *Salmonella* giant adhesin SiiE engages MUC1 and that this interaction mediates the cellular entry of *Salmonella* Enteritidis (*S*. Enteritidis) and *S*. Typhimurium. SiiE is a non-fimbrial adhesin encoded by the *Salmonella* pathogenic island 4 (SPI-4). Both SPI-4 and SPI-1, which encode the type III secretion system, are required for *Salmonella* adhesion to and invasion of polarized epithelial cells (22). We showed that neuraminidase treatment of HT29-MTX cells yields a 50% reduction in *Salmonella* invasion, suggesting a crucial role of the sialic acids on the MUC1 ED in the infection process (21).

It is not known if *Salmonella* engagement of the MUC1 ED initiates activation of downstream pathways by the MUC1 CT. In the present study, we further investigate the functions of the MUC1 ED and the contribution of the MUC1 CT domain during *Salmonella* SiiE-mediated apical invasion into intestinal epithelial cells.

## Results

### Removal of the MUC1 extracellular domain blocks *Salmonella* apical invasion into enterocytes

Human colonic HT29-MTX monolayer cultures express high levels of the transmembrane mucin MUC1 on the apical surface, and we demonstrated that *Salmonella enterica* Serovar Enteritidis can utilize MUC1 for apical invasion (21). For the current study, we set out to investigate the role of the MUC1 extracellular domain (ED) and cytoplasmic tail (CT) during *Salmonella* invasion. Confluent HT29-MTX cultures were grown for 5 days and stained with specific antibodies directed against MUC1 ED, MUC1-SEA, and MUC1 CT (Fig. 1A). All three MUC1 domains could be visualized on the apical surface of the cultures (Fig. 1B). To investigate the contribution of MUC1 ED during *Salmonella* invasion, we utilized the mucin-specific protease StcE that we previously employed to remove TM mucin O-glycosylated domains (23). HT29-MTX cultures were treated with StcE or its inactive point mutant StcE^E447D^ for 22 hours, followed by immunofluorescence staining of the different MUC1 domains. In StcE-treated cultures, the MUC1 ED was no longer detectable while the MUC1-SEA domain and MUC1 CT were still present (Fig. 1C). In StcE^E447D^-treated cultures, all three domains remained detectable. These results confirmed that StcE treatment results in the removal of the MUC1 extracellular domain while leaving the SEA domain and cytoplasmic tail intact. Next, StcE and StcE^E447D^-treated cultures were incubated with *Salmonella* for 1 hour, and bacteria were visualized by microscopy. In StcE^E447D^-treated cultures, *Salmonella* colocalized with MUC1 ED, while a reduced number of bacteria associated with StcE-treated cultures (Fig. 1D). To quantify the number of adhered/invaded bacteria, HT29-MTX wild-type and ΔMUC1 cultures were treated with StcE or StcE^E447D^, incubated with *Salmonella*-GFP for 1 hour, stained for MUC1, and analyzed by flow cytometry. Populations positive for MUC1 and/or *Salmonella* were gated into four quadrants (Q1-4, Fig. 1E, Fig. S1A). The percentage of *Salmonella*-positive cells was calculated by adding Q2 and Q3 (Fig. S1B). For StcE^E447D^-treated MUC1-WT cells, 24% of the cell population was positive for *Salmonella*. StcE treatment led to a reduction in bacterial adhesion/invasion to a similar level as the ΔMUC1 cultures (Fig. 1F). These results suggest that the MUC1 ED is essential for *Salmonella* apical invasion into enterocytes.

**Figure 1.**
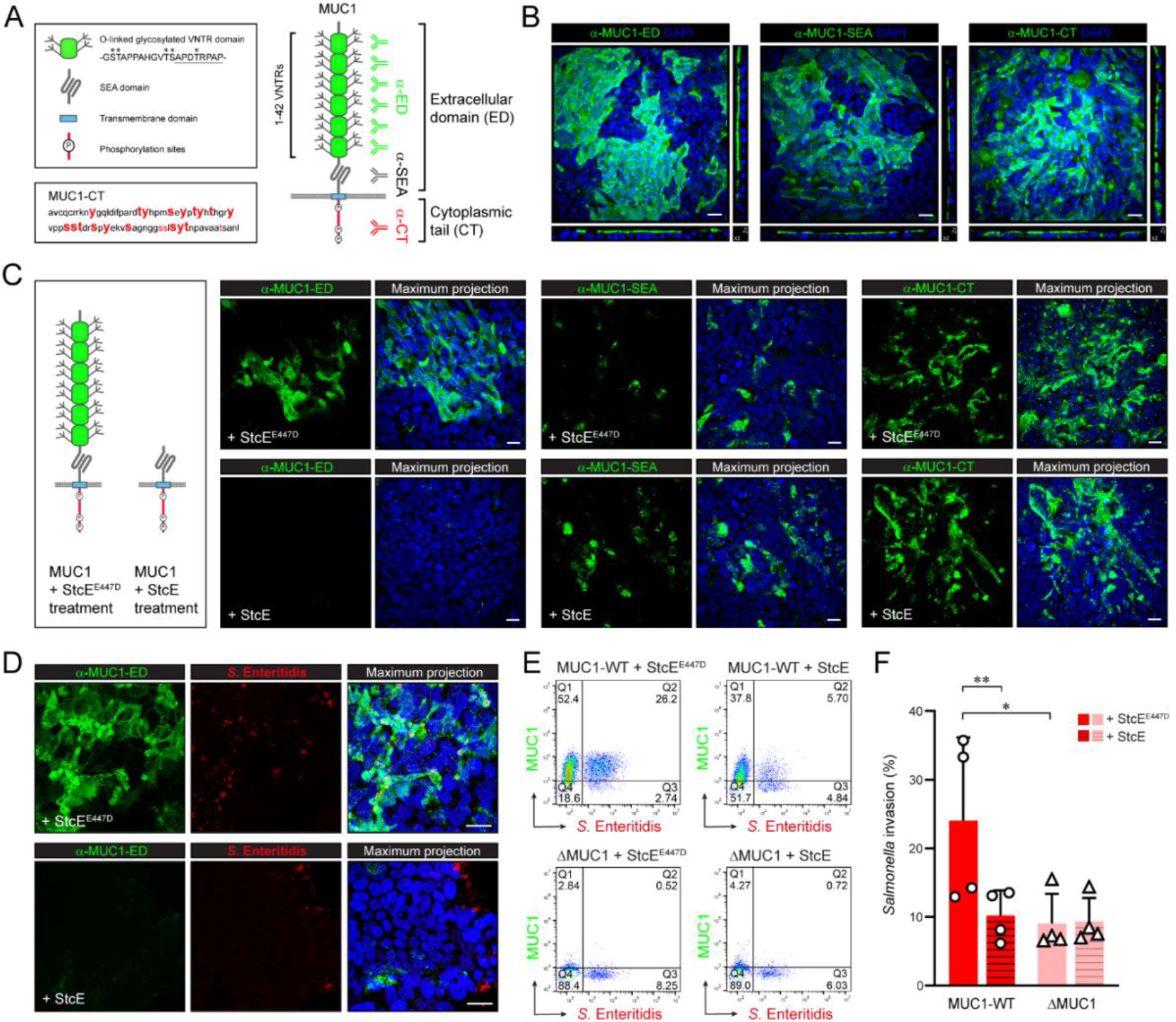
Enzymatic removal of the MUC1 extracellular domain blocks *Salmonella* apical invasion into enterocytes. A) Schematic drawing of the domain structure of transmembrane mucin MUC1 and localization of epitopes of different antibodies used in this study. B) Immunofluorescence confocal microscopy of an HT29-MTX monolayer stained with either α-MUC1-ED, α-MUC1-SEA or α-MUC1-CT2 antibodies (green) and DAPI to stain nuclei (blue). White scale bars represent 20 µm. C) Immunofluorescence confocal microscopy of HT29-MTX monolayers treated with mucin-targeting protease StcE or its inactive form StcE^E477D^ for 22 hours, stained with α-MUC1-ED antibody 139H2 (green), α-MUC1-SEA antibody 232A1 (green) or α-MUC1-CT antibody CT2 (green) in combination with DAPI to stain the nuclei (blue). White scale bars represent 20 μm. D) Immunofluorescence confocal microscopy of HT29-MTX cells treated with StcE or StcE^E477D^, infected with *Salmonella* (mCherry, red) for 1 hour, stained with α-MUC1-ED antibody 139H2 (green) and DAPI to stain the nuclei (blue). White scale bars represent 20 μm. E) Flow cytometry analysis of HT29-MTX wild-type and ΔMUC1 cultures treated with StcE or StcE^E477D^ and infected with *Salmonella* (GFP) for 1 h. After infection, monolayers were trypsinized, and single-cell solutions were fixed and stained with α-MUC1-ED antibody 139H2. F) Total percentage of *Salmonella*-positive MUC1-WT and ΔMUC1 cells (Q2+Q3) calculated from the experimental setup depicted in E. Graph depicts means ± SD of four independent experiments. Statistical analysis was performed by Student’s t-test using GraphPad Prism software. * p<0.05, ** p<0.01.

### Deletion of the MUC1 cytoplasmic tail does not impede *Salmonella* invasion

We next set out to investigate the role of the MUC1 cytoplasmic tail during *Salmonella* invasion. A targeted CRISPR/Cas9 approach was used to delete part of the MUC1 gene encoding the cytoplasmic tail while leaving the extracellular domain, SEA domain, and transmembrane domain intact. This approach resulted in a genomic deletion of about 1200 bp, and sequencing confirmed the removal of the cytoplasmic tail region (MUC1-ΔCT cells) (Fig. 2A). HT29-MTX MUC1-WT, MUC1-ΔCT, and ΔMUC1 cell lines were grown as monolayers and analyzed for expression of the different MUC1 domains. PCR, immunoblot, and immunofluorescence analysis confirmed the absence of the cytoplasmic tail in MUC1-ΔCT cultures and the absence of the full-length MUC1 protein in the ΔMUC1 cultures (Fig. 2B, C). Next, we infected MUC1-WT, MUC1-ΔCT, and ΔMUC1 cells with *Salmonella*-mCherry to determine the contribution of MUC1-CT to *Salmonella* invasion. Microscopy analysis showed that while *Salmonella* barely invaded ΔMUC1 cultures, invasion of the MUC1-ΔCT cultures was comparable to MUC1-WT (Fig. 2D). Flow cytometry analysis confirmed that *Salmonella* adhesion/invasion was comparable between MUC1-WT and MUC1-ΔCT and reduced in ΔMUC1 cultures (Fig. 2E, F). These experiments demonstrate that the MUC1 cytoplasmic tail is not essential for MUC1-mediated invasion of *Salmonella* into enterocytes.

**Figure 2:**
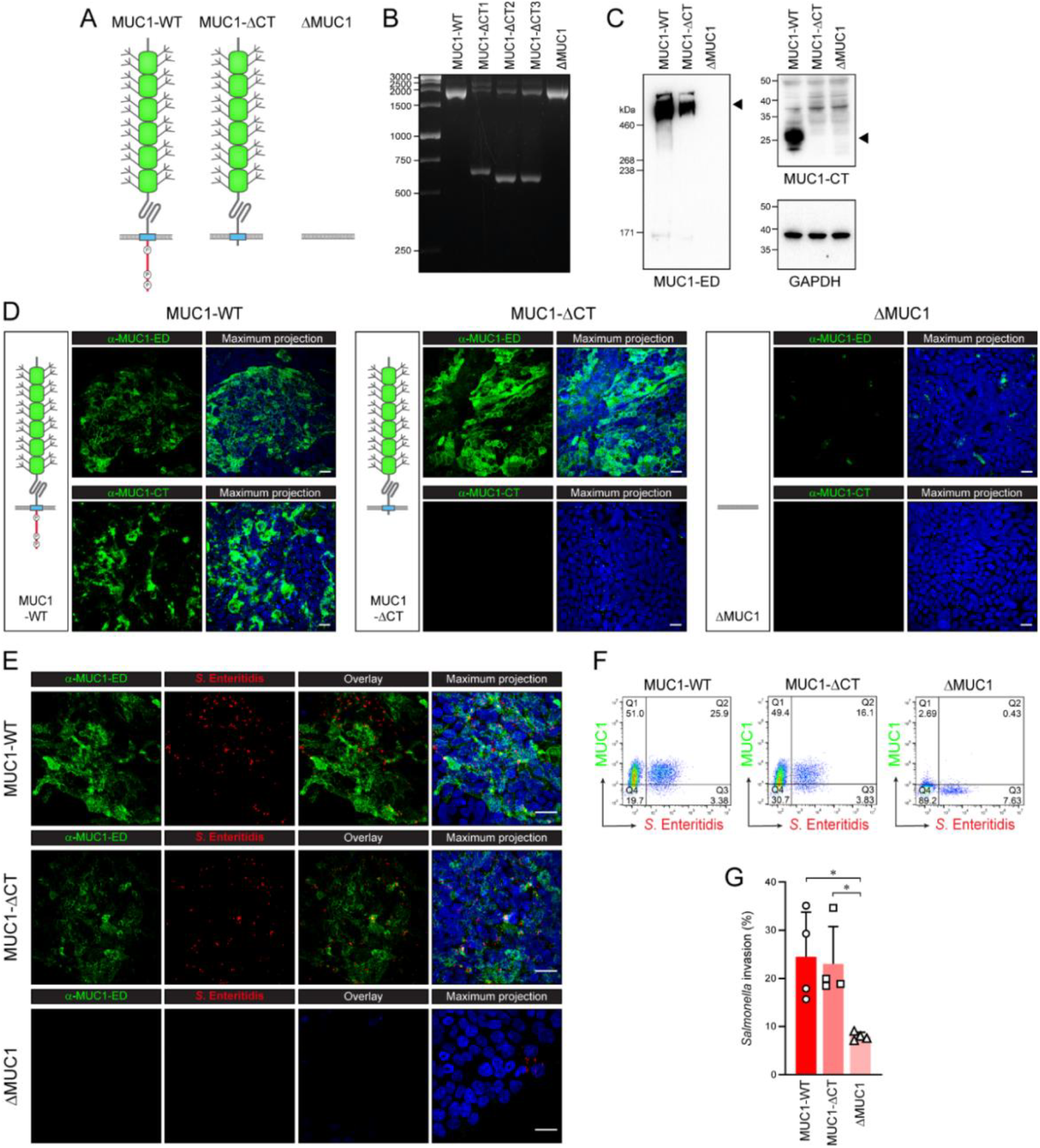
Deletion of the MUC1 cytoplasmic tail does not affect *Salmonella* invasion into enterocytes. A) Schematic drawing of MUC1 protein in wild-type cells, MUC1-ΔCT cells and ΔMUC1 cells used in this study. B) PCR-based genomic deletion analysis of CRISPR/Cas9 removal of the CT domain. Three MUC1-ΔCT clonal cell lines were confirmed to have a genomic deletion in the intended region as confirmed by sequencing. Clone MUC1-ΔCT2 was selected for subsequent experiments. C) Immunoblot analysis of HT29-MTX MUC1-WT, MUC1-ΔCT, and ΔMUC1 cultures with α-MUC1-ED (214D4) and α-MUC1-CT (CT2) and GAPDH as a loading control. D) Immunofluorescence confocal microscopy of MUC1-WT, MUC1-ΔCT and ΔMUC1 monolayers stained with α-MUC1-ED (139H2; green) or α-MUC1-CT (CT2; green) and DAPI (blue). White scale bars represent 20 μm. E) Immunofluorescence confocal microscopy of MUC1-WT, MUC1-ΔCT, and ΔMUC1 monolayers infected with *Salmonella* (mCherry, red) for 1 hour, stained with α-MUC1-ED (139H2; green) and DAPI (blue). White scale bars represent 20 μm. F) Flow cytometry analysis of MUC1-WT, MUC1-ΔCT, and ΔMUC1 cultures infected with *Salmonella* (GFP) for 1 hour. After infection, monolayers were trypsinized and single cell solutions were fixed and stained with α-MUC1-ED antibody 139H2. G) Total percentage of *Salmonella*-positive MUC1-WT, ΔMUC1, and MUC1-ΔCT cells (Q2+Q3) calculated from the experimental setup depicted in E. Graph depicts means ± SD of four independent experiments. Statistical analysis was performed by Student’s t-test using GraphPad Prism software. * p<0.05.

### Pro-inflammatory responses are similar between MUC1-WT and MUC1-ΔCT cultures

Next, we investigated the general pro-inflammatory response of MUC1-WT, MUC1-ΔCT, and ΔMUC1 cultures at 1, 3, 6, and 22 hours post *Salmonella* infection (hpi) using quantitative RT-PCR. Bacterial infection resulted in an upregulation of IL-8 and TNFα expression in all different cell types (Fig. 3A, B). At 1 hpi, IL-8 expression in ΔMUC1 cells was significantly higher compared to MUC1-WT and MUC1-ΔCT cells (Fig. 3A). At 6 hpi, both IL-8 and TNFα expression were significantly reduced in *Salmonella*-infected ΔMUC1 compared to MUC1-WT cultures, and MUC1-ΔCT cultures showed an intermediate phenotype (Fig. 3A, B). Expression of both cytokine genes was comparable at 3 and 22 hpi. Supernatants of *Salmonella*-infected cultures were collected at different time points, and IL-8 levels were assessed by ELISA. IL-8 was detectable for all three genotypes in response to *Salmonella* infection, but MUC1-WT and MUC1-ΔCT cell lines secreted significantly more IL-8 compared to ΔMUC1, which was most evident at 6 hpi (Fig. 3C, D). This result is in line with the reduced *Salmonella* invasion in the ΔMUC1 cultures compared to the MUC1-WT and MUC1-ΔCT cultures. To determine if ΔMUC1 cells have an equal capacity to produce IL-8, we stimulated MUC1-WT, MUC1-ΔCT, and ΔMUC1 cells with 1, 10, and 100 ng/mL TNFα for 6 hours. IL-8 secretion was comparable between the different genotypes at all TNFα concentrations (Fig. 3E). These results show that removal of the MUC1 cytoplasmic tail does not greatly alter gross pro-inflammatory responses upon *Salmonella* infection.

**Figure 3.**
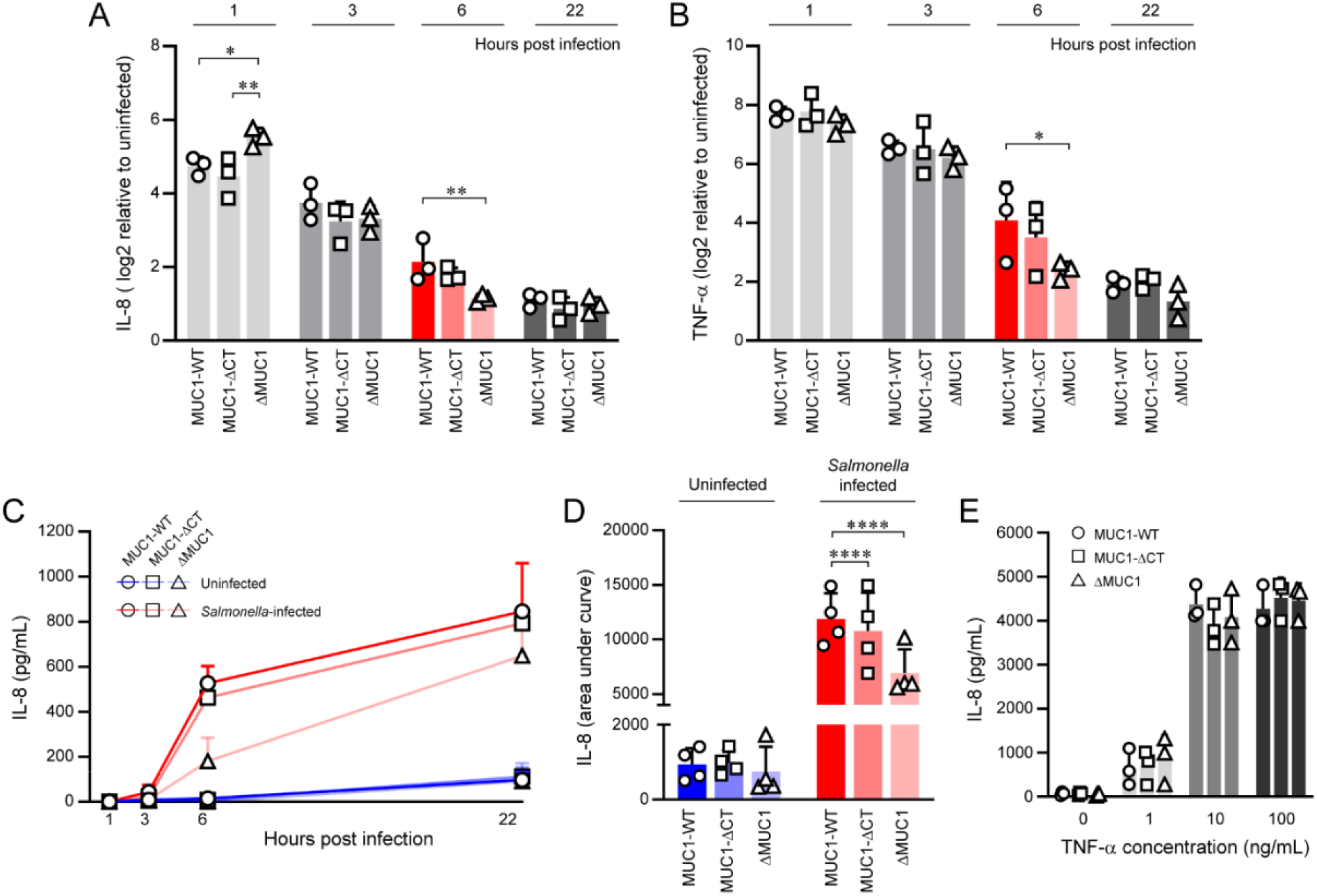
Absence of the MUC1 cytoplasmic tail does not alter major pro-inflammatory responses. A, B) Quantitative RT-PCR analysis of IL-8 and TNF-α expression in HT29-MTX MUC1-WT, MUC1-ΔCT, and ΔMUC1 cultures infected with *Salmonella* for 1h, 3h, 6h, and 22h. Plotted values are the mean log2 fold change of infected samples compared to uninfected ± SD of three independent experiments. Statistical analysis was performed by 2-way ANOVA after transformation of the values using GraphPad Prism software. * p<0.05, ** p<0.01. C) ELISA IL-8 quantification in supernatants of MUC1-WT, MUC1-ΔCT, and ΔMUC1 cultures 1, 3, 6, and 22 hours post *Salmonella* infection. Values are the mean concentration ± SD of four independent experiments. AUC values were analyzed in each experiment using GraphPad Prism software and plotted into a bar graph depicted in D. D) Area under the curve (AUC) values of IL-8 ELISA depicted in C. Statistical analysis was performed by 2-way ANOVA after transformation of the values using GraphPad Prism software. **** p<0.001. E) ELISA IL-8 quantification of supernatants of MUC1-WT, MUC1-ΔCT, and ΔMUC1 cultures treated with different (low-medium-high) concentrations of TNF-α. The graph depicts the mean concentration ± SD of three independent experiments. Statistical analysis was performed by 2-way ANOVA after transformation of the values using GraphPad Prism software.

### Transcriptomics analysis to determine contributions of MUC1-WT and MUC1-CT

To determine the contributions of the MUC1 cytoplasmic tail and full-length protein in an unbiased manner, we performed a bulk RNAseq analysis on uninfected MUC1-WT, MUC1-ΔCT, and ΔMUC1 cultures and cultures infected with *Salmonella* for 6 hours. A Principal Component Analysis (PCA) of the transcriptomes demonstrated a strong genotypic clustering between replicates and a transcriptomic switch for all genotypes upon *Salmonella* infection (Fig. 4A). Noticeably, the transcriptome of ΔMUC1 was highly distinct compared with MUC1-WT and MUC1-ΔCT transcriptomes in both uninfected and *Salmonella*-infected conditions. A direct comparison of MUC1-WT and MUC1-ΔCT transcriptomes did result in distinct clustering, pointing to a contribution of the cytoplasmic tail (Fig. 4B).

**Figure 4.**
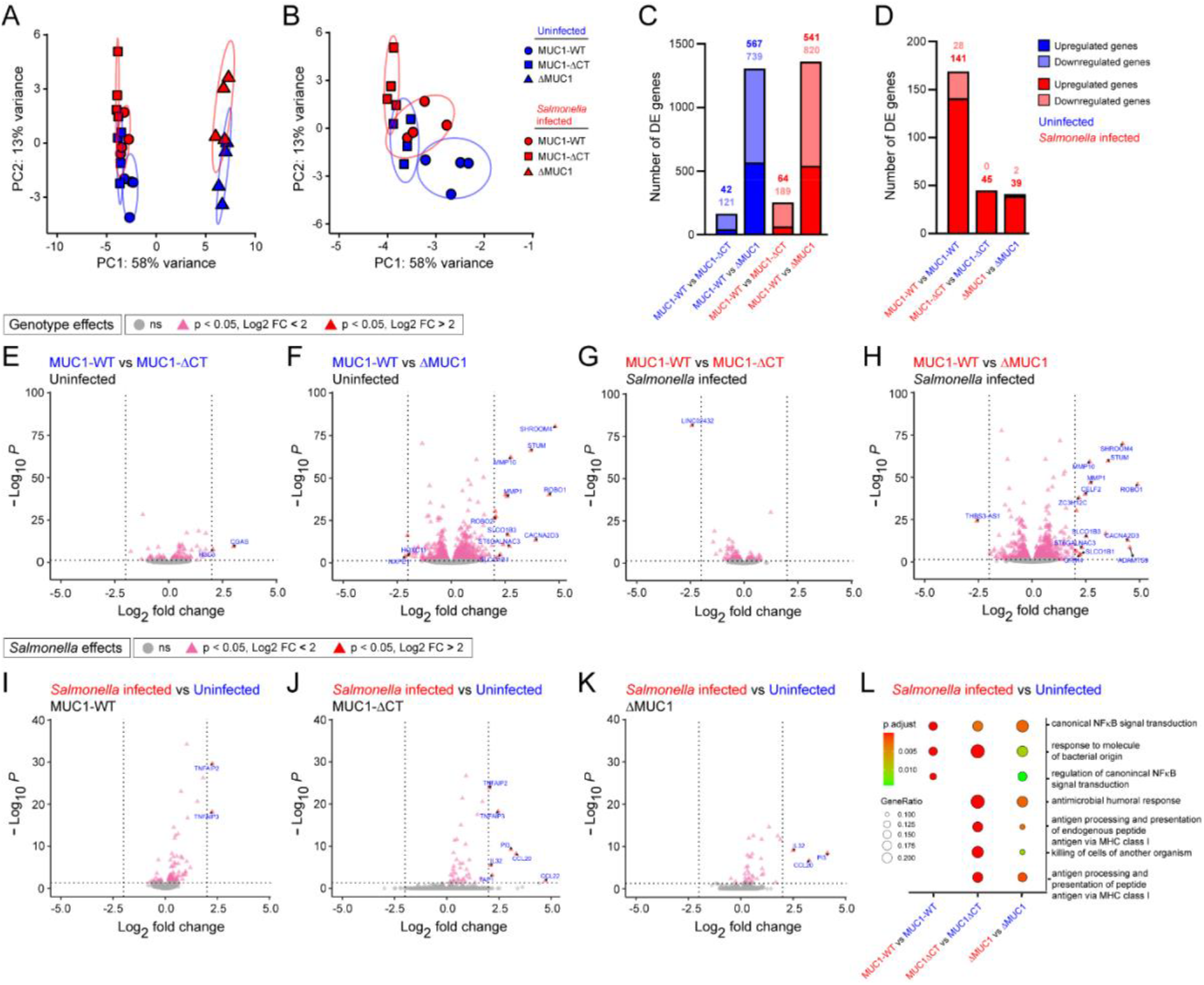
Impact of full-length MUC1 and MUC1 cytoplasmic tail on the transcriptome of uninfected and Salmonella-infected cultures. A) PCA plot of the total transcriptome of MUC1-WT (circle), MUC1-ΔCT (square), and ΔMUC1 (triangle) cells in uninfected (blue) and *Salmonella*-infected (red) conditions. B) PCA plot of the total transcriptome of MUC1-WT (circle) and MUC1-ΔCT (square) cells in uninfected (blue) and *Salmonella*-infected (red) conditions. C) Number of differentially expressed genes (DEGs) in MUC1-WT compared to MUC1-ΔCT and MUC1-WT compared to ΔMUC1 in uninfected (blue) and infected conditions (red). D) Number of DEGs in infected MUC1-WT, MUC1-ΔCT, and ΔMUC1 cells (red) compared to the respective uninfected cultures (blue). E-K) Volcano plots depicting most significantly expressed DEGs with log fold-change thresholds of −2 and 2 and p<0.05 (red) between different genotypes and between infected and uninfected samples as indicated. DEGs with adjusted p-value of <0.05 but with a lower than 2 log fold-change are depicted in pink. Non-significant genes are depicted in grey. L) Gene set enrichment analysis (GSEA) of DEGs for biological process in *Salmonella*-infected cultures of the different genotypes compared to their respective uninfected controls.

We performed differentially expressed gene (DEGs) analyses to determine the effect of genotype and *Salmonella* infection. For DEGs relating to genotype, the transcriptomes of MUC1-WT cultures were compared to ΔMUC1 and MUC1-ΔCT cultures in uninfected and *Salmonella*-infected conditions. Comparison of MUC1-WT with ΔMUC1 resulted in 1306 (uninfected) and 1361 (infected) DEGs. Comparison of MUC1-WT with MUC1-ΔCT led to 163 (uninfected) and 253 (infected) DEGs (Fig. 4C, supplementary tables 1-4). For DEGs relating to *Salmonella* infection, the transcriptome of a *Salmonella*-infected genotype was compared to its respective uninfected control, resulting in 169 (MUC1-WT), 45 (MUC1-ΔCT), and 41 (ΔMUC1) DEGs (Fig. 4D, supplementary tables 5-7). The main findings of the DEG analysis are that there is a strong genotypic effect in ΔMUC1 cells and that MUC1-WT cultures differentially express a larger set of genes upon *Salmonella* infection compared to MUC1-ΔCT and ΔMUC1 cultures.

We further analyzed the most significant alterations relating to genotype by plotting the DEGs in volcano plots and highlighted those with changes of log2 fold −2 and 2 and adjusted p-value <0.05 (Fig. 4E-H). This analysis again pointed to larger transcriptional changes linked to the ΔMUC1 genotype (Fig. 4F, H) compared to the MUC1-ΔCT genotype (Fig. 4E, G). The most significant alterations relating to *Salmonella* infection were also plotted in volcano plots with the same thresholds (Fig. 4I-K). A limited number of genes were differentially expressed more than 2-fold. In MUC1-WT cells, the genes TNFAIP2 and TNFAIP3 were the most highly induced. In MUC1-ΔCT cells, TNFAIP2, TNFAIP3, TAP1, PI3, IL32, CCL20, and CCL22 were the most significantly upregulated genes and in ΔMUC1 cultures PI3, IL32, and CCL20. The DEGs from our transcriptomics dataset were also used for gene set enrichment analysis (GSEA) of the biological processes. Overall, the analysis demonstrated enrichment of gene clusters associated with processes including defense response to bacteria and inflammatory response across all three cell cultures (Fig. 4L). MUC1-WT cultures displayed a distinct enrichment profile compared to MUC1-ΔCT and ΔMUC1 cultures. Processes related to antigen processing and immune responses were enriched in MUC1-ΔCT and ΔMUC1 cultures. In MUC1-WT and not in the other genotypes, a cluster of DEGs related to canonical NFκB signaling transduction was significant. This suggests that the MUC1 cytoplasmic tail (CT), or the full-length protein, might be involved in the NFκB signaling.

### An NFκB-related gene cluster is uniquely associated with *Salmonella* infection in MUC1-WT

To better understand the characteristics of the transcriptional responses found in the different genotypes, we determined the DEGs found in infected vs uninfected cultures in the different genotypes (Fig. 5A, B). The upregulation of 24 genes was shared between the three genotypes, and most of these genes were immune genes related to infection (Fig. 5C). 132 genes were differentially expressed upon infection in the MUC1-WT cultures but not in MUC1-ΔCT or ΔMUC1 cultures (Fig. 5A, B). Of this subset, 104 genes were up-regulated and 28 genes down-regulated. A heatmap analysis of the 132 MUC1-WT associated genes demonstrated a clear cluster of infection-upregulated genes that was absent in MUC1-ΔCT and ΔMUC1 cultures (Fig. 5D). Among this cluster were again several genes relating to the NFκB pathway, and GSEA analysis of biological processes also demonstrated a significant association with NFκB signaling transduction (Fig. 5E). Next, we used the ChEA3 database (24) to identify transcription factors potentially involved in the regulation of the 132 DEGs associated with *Salmonella* infection in MUC1-WT cultures. Top hits of the ChEA3 analysis included RELB and NFKB2, which are main transcription factors of the non-canonical NFκB pathway (Fig. 5F). Genes related to RELB and NFKB2 were plotted in a heatmap to visualize their expression levels in non-infected and infected cultures. Again, a cluster of genes unique to MUC1-WT responses could be observed. In conclusion, our RNAseq analysis points towards involvement of the MUC1 cytoplasmic tail (and full-length protein) in the NFκB signaling pathway upon *Salmonella* infection.

**Figure 5.**
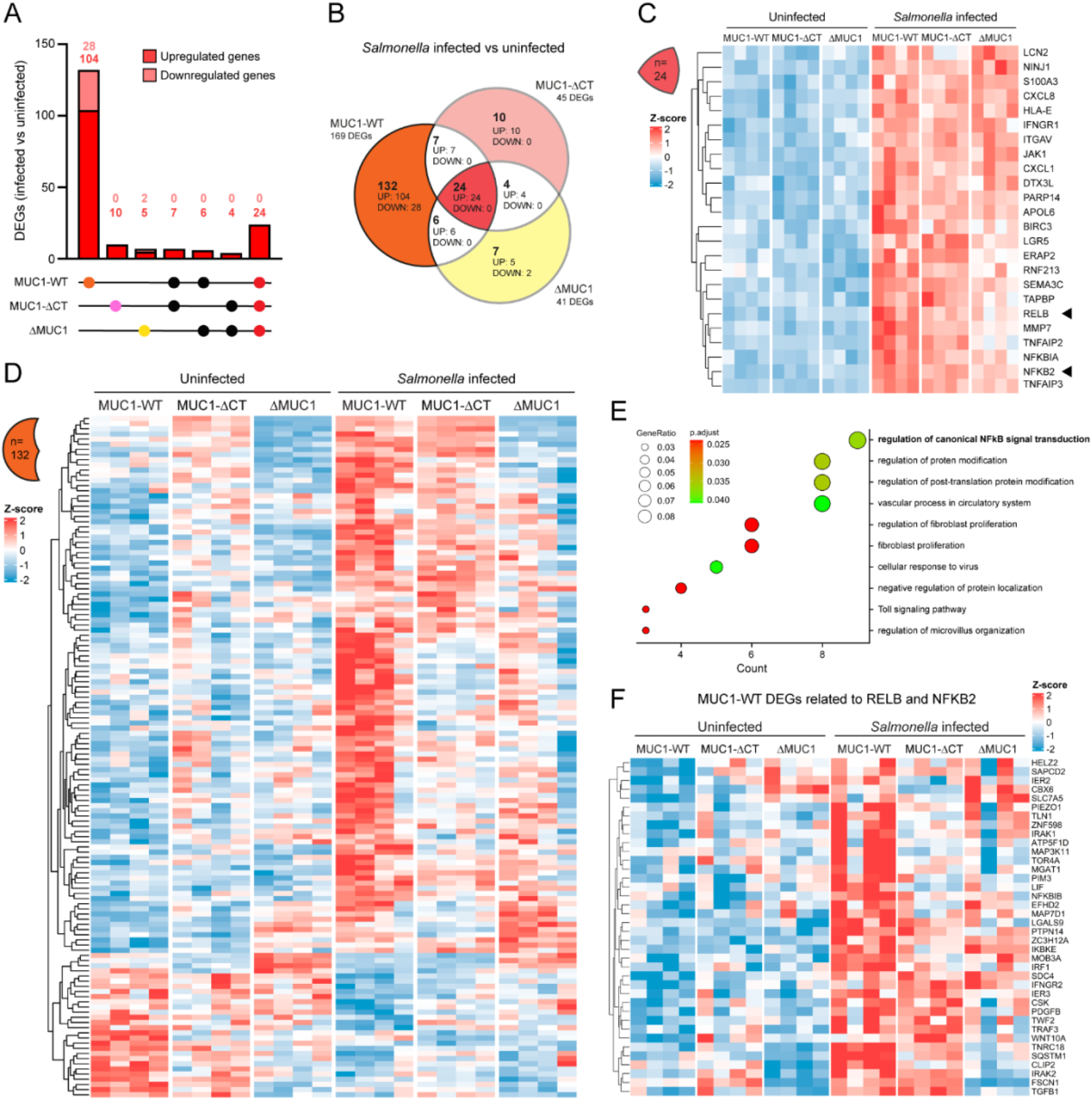
*Salmonella* infection results in upregulation of NFκB-related DEG cluster in MUC1-WT but not MUC1-ΔCT and ΔMUC1 cells. A) Number of DEGs in each cell line comparing *Salmonella*-infected cultures to their respective uninfected controls. B) Venn diagram of DEGs found in MUC1-WT, MUC1-ΔCT, and ΔMUC1 when comparing *Salmonella*-infected cultures to uninfected cultures. C) Heatmap displaying the 24 shared DEGs induced during *Salmonella* infection in the three genotypes. D) Heatmap displaying the 132 DEGs uniquely induced in MUC1-WT cultures but not MUC1-ΔCT and ΔMUC1 cultures during *Salmonella* infection. E) GSEA for the biological process using the profileR database of 132 DEGs uniquely found in MUC1-WT during *Salmonella* infection. F) Heatmap displaying DEGs related to the NFKB2 and RELB signaling pathway uniquely found in *Salmonella*-infected MUC1-WT cultures.

### Deletion of MUC1 or its cytoplasmic tail alters expression of NFκB proteins

NFκB signaling can broadly be divided into two distinct pathways. In the canonical NFκB pathway, RelA is the transcriptional activator that is regulated by Nfkb1. The non-canonical NFκB pathway consists of transcriptional activator RelB and regulator NfkB2 (Fig. 6A). To determine the contributions of the different MUC1 domains to NFκB signaling, we quantified the levels of NFκB proteins in lysates of uninfected and *Salmonella*-infected MUC1-WT, MUC1-ΔCT, and ΔMUC1 cultures after 6 hours of infection.

**Figure 6.**
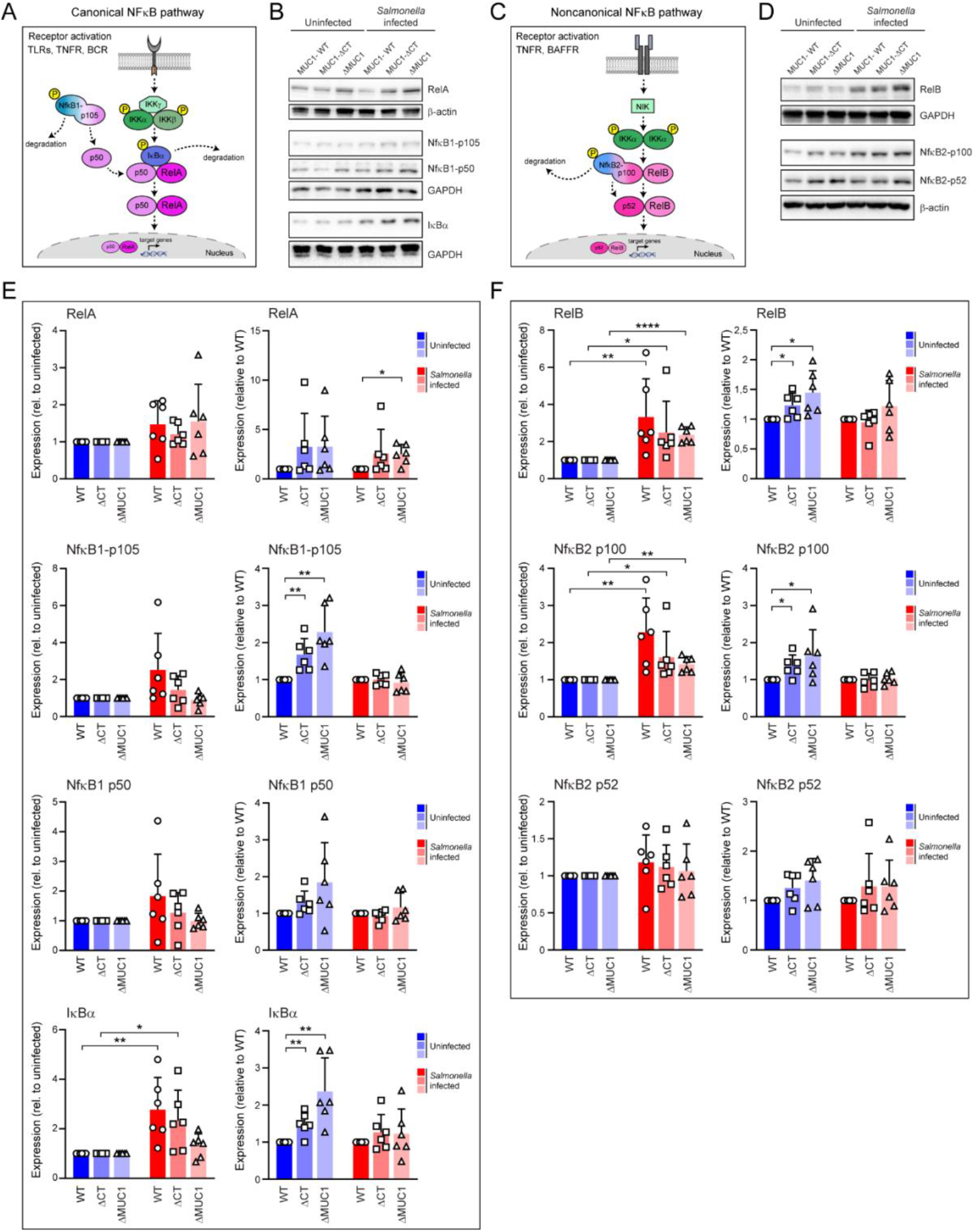
Expression of RelB of the non-canonical NFκB signaling pathway is altered in MUC1-ΔCT and ΔMUC1 cells. A) Schematic of the activation of the RelA inhibitory subunits and transcriptional subunit in the canonical NFκB signaling pathway. B) Representative western blots of MUC1-WT, MUC1-ΔCT, and ΔMUC1 cells infected with *Salmonella* for 6 hours, blotted for canonical NFκB signaling pathway: RelA, NfκB1-p105 and p50, and IκBα. GAPDH or β-actin was used as a loading control. C) Schematic of the activation of the RelB inhibitory subunits and transcriptional subunit in the non-canonical NFκB signaling pathway. D) Representative western blots of MUC1-WT, MUC1-ΔCT, and ΔMUC1 cells infected with *Salmonella* for 6 hours, blotted for non-canonical NFκB signaling pathways: RelB and NfκB2-p100 and p52. GAPDH or β-actin was used as a loading control. E) Quantification of RelA, NfκB1-p105 and p50, and IκBα expression by western blot as depicted in (B). Graphs depict relative expression in infected cells relative to uninfected controls (left) and relative expression to MUC1-WT in uninfected and infected cells (right). Values are the mean ± SD of six independent experiments. Statistical analysis was performed by a one-sample t-test using GraphPad Prism software. F) Quantification of RelB and NfκB2-p100 and p52 expression by western blot as depicted in (D). Graphs depict relative expression in infected cells relative to uninfected controls (left) and relative expression to MUC1-WT in uninfected and infected cells (right). Plotted values are the mean ± SD of six independent experiments. Statistical analysis was performed by a one-sample t-test using GraphPad Prism software. * p<0.05, ** p<0.01, **** p<0.0001.

For the canonical pathway, we determined expression of the transcriptional activator RelA, the NFκB1 precursor (NFκB1-p105), its processed transcriptional product (NFκB1-p50), and the inhibitory subunit IκBα by immunoblot (Fig. 6B, D). RelA protein levels were comparable between MUC1-WT, MUC1-ΔCT, and ΔMUC1 cultures following *Salmonella* infection relative to their respective uninfected controls (Fig. 6D, left). Under uninfected conditions, RelA expression in MUC1-ΔCT and ΔMUC1 cells did not significantly differ from that in MUC1-WT cells (Fig. 6D, right). Following infection, induction of RelA expression was significantly higher in ΔMUC1 cells compared to MUC1-WT (Fig. 6D, right). In uninfected cultures, NFκB1-p105 levels were significantly higher in MUC1-ΔCT and ΔMUC1 cells relative to MUC1-WT, suggesting altered regulation of the precursor form. This increase was not observed for NFκB1-p50 (Fig. 6D, right). Following *Salmonella* infection, IκBα levels were significantly upregulated in MUC1-WT and MUC1-ΔCT but not in ΔMUC1 cultures (Fig. 6D, left). Under uninfected conditions, IκBα expression was higher in MUC1-ΔCT and ΔMUC1 cells compared to MUC1-WT.

For the non-canonical pathway, we determined expression of the transcriptional activator RelB, NFκB2 precursor (NFκB2-p100), and its processed transcriptional product (NFκB2-p52) (Fig. 6C, E). Upon *Salmonella* infection, RelB and NFκB2-p100 levels were significantly induced in all three cell types compared to the uninfected cultures (Fig. 6E, left). However, expression of the transcriptionally active NFκB2-p52 subunit was comparable between conditions. Under uninfected conditions, RelB and NFκB2-p100 levels are higher in MUC1-ΔCT and ΔMUC1 cells compared to MUC1-WT cells, potentially indicating a role for the MUC1 cytoplasmic tail. Again, no changes were observed for the NFκB2-p52 transcriptional subunit.

In conclusion, we observed that under uninfected conditions, the presence of MUC1-WT or the MUC1 cytoplasmic tail results in lower expression of several components of the canonical and non-canonical NFκB pathway compared to the absence of MUC1 CT. RelB, the transcriptional activator of the non-canonical pathway, is most strongly induced upon *Salmonella* infection, which is in line with the RNAseq analysis and the most relevant gene cluster associated with MUC1-WT-associated responses.

### Functional consequences of altered MUC1-mediated signaling for cytokine release

In a final set of experiments, we examined the contributions of the full-length MUC1 and MUC1-CT to the secretion of cytokines and immune factors during *Salmonella* infection. A Luminex assay was used to determine the presence of selected pro-inflammatory immune analytes in supernatants of uninfected and *Salmonella*-infected MUC1-WT, MUC1-ΔCT, and ΔMUC1 cultures. In line with our RNAseq and immunoblot data, we observed that MUC1-WT cultures displayed an overall stronger inflammatory immune response compared to MUC1-ΔCT and ΔMUC1 cultures after *Salmonella* invasion (Fig. 7A). In the ΔMUC1 cultures, in which *Salmonella* invasion is severely reduced (Figs. 1,2), none of the selected analytes were significantly upregulated upon infection (Fig. 7B, C). These results demonstrate that in this intestinal model, *Salmonella* invasion through MUC1 is absolutely required to induce secretion of the immune factors in our panel. IL-8 was the most strongly induced cytokine upon *Salmonella* infection of MUC1-WT and MUC1-ΔCT cultures and also MIP1α, MCP1, IL-18 and TNF-α were significantly induced upon infection. In infected MUC1-ΔCT cultures, IL-18 secretion was significantly reduced compared to MUC1-WT (Fig. 7B). Among the secreted immune factors, galectin-3 and galectin-9 were most highly induced by *Salmonella* infection. In infected MUC1-ΔCT cultures, secretion of galectin-3 was significantly lower compared to MUC1-WT and a similar trend, although not significant, was observed for galectin-9. sICAM, S100A8 and PDGFBB were also detectable in the supernatants of infected MUC1-WT and MUC1-ΔCT cultures, but no significant differences were observed relating to the absence of the CT (Fig. 7C). We conclude that the MUC1 invasion pathway is essential for the induction of secretion of cytokines and immune factors. The MUC1 CT is essential for optimal immune factor secretion and specifically IL-18 and galectin-3 are significantly reduced in the supernatants of infected MUC1-ΔCT cultures. Taken together, our results demonstrate that the MUC1 ED is essential for *Salmonella* invasion and that the MUC1 CT plays a role in modulating the NFκB pathway and mounting a full pro-inflammatory response during infection.

**Figure 7.**
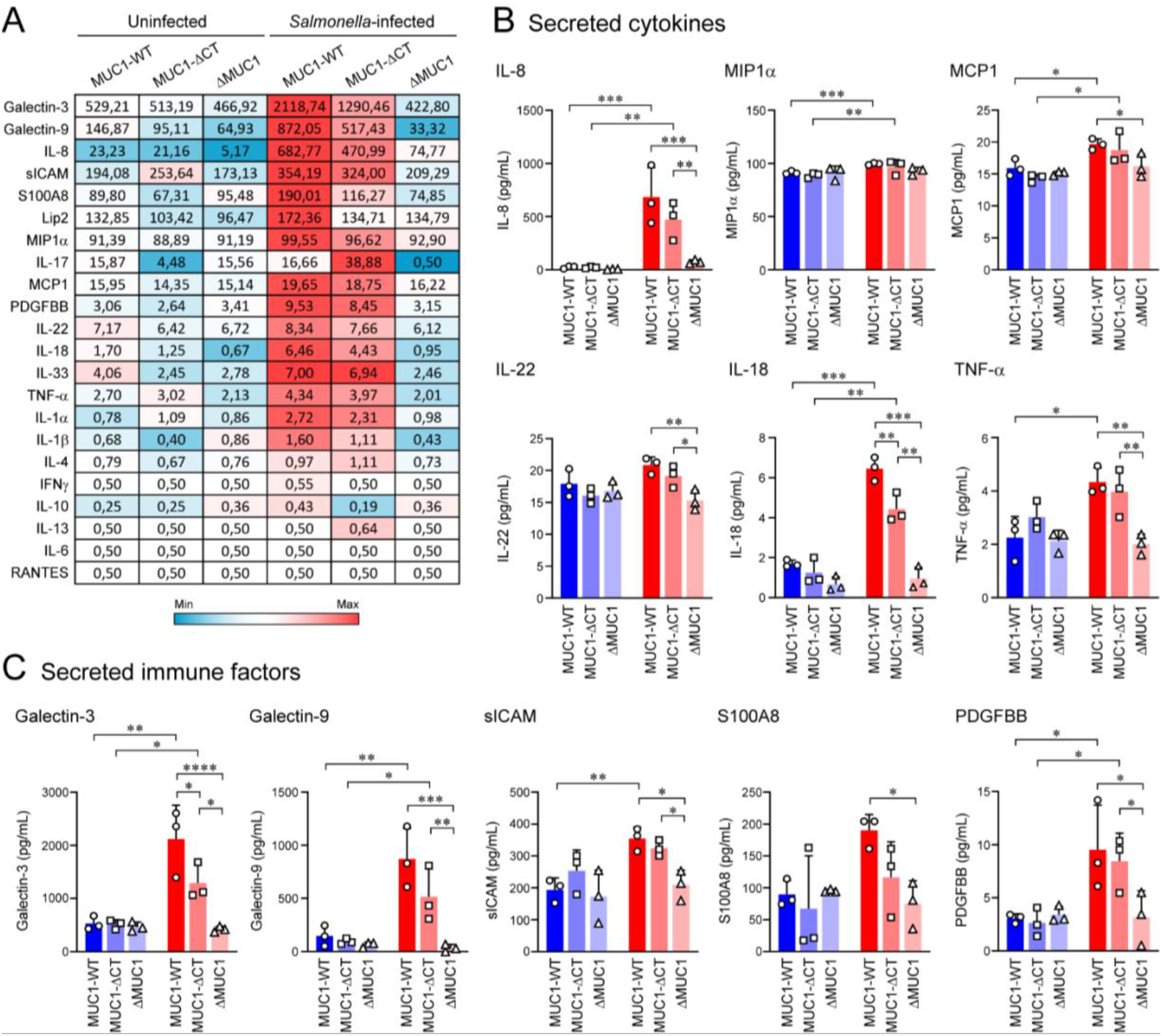
Functional impact of MUC1-mediated NFκB alterations on cytokine profiles in uninfected and *Salmonella*-infected cultures. A) Heatmap of all Luminex analyses of supernatants of MUC1-WT, MUC1-ΔCT, and ΔMUC1 cultures infected with *Salmonella* for 6 hours. Values are the mean concentration in pg/mL of three independent experiments. The range for the heatmap gradient for each analyte was set from 0 to the highest concentration detected per analyte. B) Graphs depicting Luminex data of secreted cytokines panel: IL-8, MIPα, MCP1, IL-22, IL-18 and TNF-α. Values are the mean concentration in pg/mL ± SD of three independent experiments. C) Graphs depicting Luminex data of other secreted immune factors detected in this panel: galectin-3, galectin-9, sICAM, S100A8, and PDGFBB. Plotted values are the mean concentrations in pg/mL ± SD of three independent experiments. Statistical analysis was performed by 2-way paired ANOVA using GraphPad Prism software. * p<0.05, ** p <0.01,*** p<0.001, ****p<0.0001.

## Discussion

We previously demonstrated that *Salmonella* expresses an adhesin called SiiE that can engage the transmembrane mucin MUC1 for apical invasion into enterocytes (21). In the current study, we found that the glycosylated MUC1 ED but not the MUC1 CT is essential for MUC1-mediated *Salmonella* invasion and major pro-inflammatory response (Fig. 1,2,3). Removal of the full MUC1 protein (ΔMUC1) greatly altered gene expression, while deletion of the cytoplasmic tail (MUC1-ΔCT) only had a moderate effect on the transcriptome. A gene cluster relating to the non-canonical NFκB pathway was uniquely induced in wild-type cells and not in MUC1-ΔCT or ΔMUC1 upon infection (Fig. 4,5). In uninfected cultures, the MUC1 protein and its cytoplasmic tail played a role in suppressing NFκB components (Fig. 6). During *Salmonella* infection, the presence of the full-length MUC1 protein was required to mount the strongest inflammatory response (Fig. 7). Our results demonstrate that the extracellular domain and cytoplasmic tail of MUC1 play unique roles during *Salmonella* invasion and infection.

MUC1 is thought to play a central role in many bacterial and viral interactions at mucosal surfaces. It interacts with epithelial receptors such as TLRs, EGFR, and B1 integrin (14, 25) and intracellular signaling pathways. The extracellular domain can be adhesive or have anti-adhesive (barrier) properties. Shedding of the ED from the SEA domain or the plasma membrane could serve as a decoy to protect the epithelium. MUC1 protects against *Campylobacter jejuni* and *Helicobacter pylori* infection (26, 27). During some microbe-host interactions, MUC1 appears to exert overall anti-inflammatory effects (26). However, this effect seems to be context-dependent, as MUC1 was also shown to contribute to proinflammatory responses in tumor cells (28, 29). Due to the pleotropic functions of the different MUC1 domains, the underlying mechanisms are difficult to resolve.

Glycosylated EDs of transmembrane mucins extend into the lumen of the intestine and are, therefore, naturally the point of contact for both commensal bacteria and invading pathogens. We confirmed the importance of the MUC1 ED for *Salmonella* invasion (Fig. 1) and this is in line with our previous finding that *Salmonella* SiiE binding to MUC1 was dependent on sialic acids that are often the terminal sugars of O-glycan structures (21). The importance of the MUC1 extracellular domain has also been established for other bacteria (30). For example, adhesion of *H. pylori* to gastric epithelial cells was shown to be dependent on the MUC1-VNTR domain (31). Polymorphisms that affect the MUC1 VNTR are linked to increased susceptibility to *H. pylori* infection in the stomach (32, 33). *Pseudomonas aeruginosa* interacts with the MUC1 ectodomain on the surface of respiratory cells or in its shed form (34). Interestingly, in our study, deletion of the full-length MUC1, including the glycosylated ED, had a great impact on the total transcriptome, also in non-infected cultures (Fig. 4). We did not address these changes in detail because *Salmonella* invasion was reduced due to the absence of the MUC1 entry receptor. However, this dataset can form the basis for future studies into the diverse functions of MUC1. Another striking finding was that in our intestinal HT29-MTX model *Salmonella* apical invasion through the MUC1 pathway was essential for secretion of all the selected pro-inflammatory cytokines and immune factors (Fig. 7). As infection of ΔMUC1 cultures did not induce secretion of any immune factor, we conclude that invasion through the SiiE-MUC1 route is absolutely required for mounting an immune response in this intestinal model.

The MUC1 CT consists of 72 amino acids and contains multiple putative phosphorylation sites, suggesting signaling potential (11, 35). MUC1 is overexpressed in different types of adenocarcinomas and has been associated with enhanced cell proliferation (36, 37), altered cell–cell interactions (38), and increased metastasis (39, 40). Some of these effects are (in part) mediated by the CT, for example, through interaction with β-catenin (41, 42), GSK3β (43) c-Src (44), and PKC (45). It is not clear if MUC1 CT also exerts these functions in the healthy intestinal epithelium or during interactions with microbes. Our experiments demonstrate that the MUC1 CT is not essential for MUC1-mediated *Salmonella* invasion (Fig. 2) but that the CT contributes to regulation of NFκB responses and secretion of pro-inflammatory immune factors (Fig. 3-6). A distinct cluster of 132 genes significantly associated with transcription factors RelB and NFKB2 was differentially regulated in MUC1-WT cultures upon *Salmonella* infection but not in MUC1-ΔCT cultures (Fig. 5). The more robust transcriptional responses of MUC1-WT cells compared to MUC1-ΔCT cells was in line with a previous study in which overexpression of a full-length MUC1 construct led to stronger transcriptional responses in pancreatic tumor cells compared to a MUC1-ΔCT construct (46). It should be noted that many studies make use of MUC1 overexpression to mimic cancer status, which may alter signaling behavior compared to endogenous expression levels. In our study, we are studying endogenous expression levels of MUC1, but the colorectal carcinoma background of the HT29-MTX cells might also affect MUC1 signaling.

Our unbiased RNAseq approach demonstrates that regulation of the NFκB pathway and its downstream effectors is the most prominent function of the MUC1 CT in *Salmonella*-infected HT29-MTX cells. The link between MUC1 CT and the NFκB is in line with several published works. In *H. pylori-infected* human gastric epithelial cells, MUC1 CT was shown to interact with IKKγ, the inhibitory component of the canonical NFκB pathway (26). During infection, MUC1 suppressed NFκB activation and IL-8 production, and this effect seems to be dependent on IKKβ (26). Overexpression of MUC1 or a chimeric protein consisting of the extracellular domain of CD8 and the CT of MUC1 (pCD8-MUC1) reduced NFκB activation, and IKKγ could be co-immunoprecipitated (26). In tumor cells, an interaction between MUC1 CT and the RelA (NFκB-p65) transcription factor of the canonical pathway was demonstrated, and this interaction contributes to pro-inflammatory responses (26, 28, 29). MUC1 can also partner with TLRs such as TLR5, and this interaction reduces MyD88 recruitment and dampens NFκB signaling pathways (14, 16, 47). *In vivo* experiments demonstrated that Muc1 suppressed NFκB and NLRP3 inflammasome activation and IL-1β responses in mice subjected to chronic *H. pylori* infection (47). Our data demonstrated that several components of the canonical and non-canonical NFκB pathway were differentially expressed in MUC1-WT compared to MUC1-ΔCT and ΔMUC1 cultures. Under uninfected conditions, MUC1-WT cells had lower expression of both NFκB1 p105 and NFκB2 p100 regulatory units (Fig. 6). This difference in expression was not observed in *Salmonella*-infected cultures, perhaps because the strong infection stimulus overrode the modulatory effect of MUC1. Notably, RelB and NFκB p100, and not RelA and NFκB p105, were markedly induced upon *Salmonella* infection in all cell types, indicating a predominant activation of the non-canonical pathway in this model. This is surprising as the canonical NFκB pathway is normally considered to be the main branch triggered by bacterial invasion. The non-canonical pathway has been described to remain active during extended infections and support the survival of host cells (48, 49). In the intestine, crosstalk between both branches of the NFκB pathway is required to initiate optimal innate immune responses to pathogenic stimuli (50). Our findings reveal an interesting link between MUC1 and the non-canonical NFκB pathway, which should be taken into consideration in future studies.

Functionally, the reduced *Salmonella* invasion in ΔMUC1 cultures and altered regulation of the NFκB pathway in MUC1-ΔCT leads to lowered immune responses upon *Salmonella* infection. The contribution of the CT is moderate, but this domain is necessary for maximum induction of cytokine and immune factor secretion during infection. Interestingly, secretion of galectin-3 and IL-18 was significantly reduced in the absence of the CT (Fig. 7). Galectin-3 and galectin-9 are beta-galactosidase-binding lectins that have different immune-modulatory functions. An important function of galectin-3 is its interaction with bacterial lipopolysaccharide (LPS) which induces phagocytic killing (51) and protects against invasion into the mucus layer (52). Galectin-3 can also directly interact with the MUC1 ED leading to mucin clustering and exposure of underlying surface receptors (53, 54). IL-18 is a pro-inflammatory cytokine that induces interferon-gamma, regulatory T cells and NK cells (55, 56) and has not previously been linked to MUC1 downstream signaling. Further research is necessary to understand the function of MUC1 in regulation of galectin and IL-18 responses. The specific functions of the MUC1 ED and CT during *Salmonella* invasion and subsequent host responses warrant further investigations into the functions of different TM mucins at the intestinal microbe-host interface.

## Materials and Methods

### Bacterial culture

*Salmonella enterica* serovar Enteritidis (*S*. Enteritidis) strain 90-13-706 (CVI, Lelystad) was used for all *Salmonella* experiments. An *S*. Enteritidis strain transformed with an mCherry plasmid was previously described (21). To generate a GFP-expressing strain, *S*. Enteritidis was transformed with plasmid pMW85 expressing GFP from a PpagC promoter, generously provided by Dr. Dirk Bumann (57). All *Salmonella* strains were routinely cultured on LB agar plates and LB broth containing ampicillin (100 μg/ml) or kanamycin (50 μl/ml) at 37°C.

### Cell culture

The human colorectal carcinoma cell lines HT29-MTX (MUC1-WT), HT29-MTX-ΔMUC1 (ΔMUC1)(21), and HT29-MTX-MUC1-ΔCT (MUC1-ΔCT) (this study) were routinely cultured in 25 cm^2^ flasks in Dulbecco’s modified Eagle’s medium (DMEM) supplemented with 10% fetal calf serum (FCS) at 37°C in 10% CO_2_. For flow cytometry, RNA sequencing, western blots, ELISA, and Luminex experiments, 500,000 cells or cells at 50% confluency were seeded into 12-well plates and allowed to grow to full confluency and differentiate for 6 days. For imaging experiments, 250,000 cells or cells at 50% confluency were seeded on circular glass coverslips (8 mm diameter #1.5) in 24-well plates and grown for 6 days. StcE mucinase and the enzymatically inactive StcE^E447D^ were expressed and purified as described previously (25). For mucinase treatment, confluent cultures in 12-well plates (about 3 million cells per well) were treated with 15 µg of StcE or StcE^E447D^ in 1 ml of DMEM with 10% FCS for 20 hours at 37°C, 10% CO_2_. After treatment, cells were washed with DPBS or DMEM without FCS before infection assay experiments.

### Targeted deletion of the MUC1 CT using CRISPR/Cas9

We designed a targeted CRISPR/Cas9 strategy to specifically delete the MUC1 cytoplasmic tail in HT29-MTX cells while retaining the extracellular and transmembrane domains,. The pCRISPR-hCas9-2xgRNA-Puro plasmid that encodes Cas9 with 2 gRNAs was used to generate a 1235 bp deletion at the 3’ end of the MUC1 gene. The pCRISPR plasmid was digested with SapI and simultaneously dephosphorylated with alkaline phosphatase (FastAP; ThermoFisher). gRNA primer sets A (5’-GCTGCCCGTAGTTCTTTCGG-3’ and 5’-CCGAAAGAACTACGGGCAGC-3’) and B (5’-GGCGACGTGCCCCTACAAGT-3’ and 5’-ACTTGTAGGGGCACGTCGCC-3’) were phosphorylated with T4 polynucleotide kinase (ThermoFisher) at 37°C for 30 minutes and annealed by cooling down from 85°C to 25°C at 0.1°C/sec. Annealed primer sets were ligated into the SapI-digested pCRISPR plasmid as confirmed by sequencing with primers KS46 5’-GTTCACGTAGTGCCAAGGTCG-3’ and KS47 5’-GAGTCAGTGAGCGAGGAAGC-3’, resulting in plasmid pCRISPR. HT29-MTX cells were grown in a 25cm^2^ flask, trypsinized, and transfected in suspension with either pCRISPR, pCRISPR-empty, or no plasmid using Fugene (Promega) with a Fugene: plasmid ratio of 3:1. After culturing of 2 days in DMEM supplemented with 10% FCS, 5 μg/ml puromycin (Life Technologies) was added in suspension and cells were grown for 5 days to kill undelivered cells. Once all negative control cells had died, single cells were selected by making serial dilutions, and single-cell clones were tested for the MUC1 CT deletion by PCR with primers KS304 5’-CTCTCGATATAACCTGACGATCTCAGACG-3’ and KS305 5’-CACACACTGAGAAGTGTCCGAGAAATTG-3’, sequencing, immunofluorescence microscopy, and by western blot.

### Infection assays

*Salmonella* was grown overnight on LB agar (Biotrading), and the next day, a single colony was picked and inoculated in 10 mL of LB medium (Biotrading) containing ampicillin (100 μg/ml) or kanamycin (50 μl/ml). and incubated overnight at 37°C while shaking at 160 rpm. To induce optimal SiiE expression, the overnight *Salmonella* culture was inoculated at 1:1000 dilution in 25 mL LB medium without antibiotics and incubated at 37°C while shaking at 160 rpm and grown till late logarithmic phase to an OD_600_ of 1. 1 ml of bacterial culture was centrifuged for 2 minutes at 8000 rpm and the pellet was resuspended in 1 ml DPBS. Quickly after making the bacterial suspension, bacteria were added to the confluent HT29-MTX cultures grown in 12-well or 24-well plates in DMEM without FCS. <u>For flow cytometry</u>, bacteria were added at an MOI of 15 to the intestinal cultures. Monolayers were washed twice with DPBS and trypsinized with 200 µL of trypsin for 5 minutes at 37°C at 10% CO_2_. Cells were taken up in cold FACS buffer (2% BSA in DPBS) and centrifuged at 1,500 rpm for 5 minutes. Cells were fixed with 4% cold paraformaldehyde in PBS for 10 minutes. <u>For fluorescence microscopy</u>, bacteria were added at an MOI of 15 to the intestinal cultures grown on glass slides. After incubation with bacteria, cultures were washed twice with DPBS and fixed with 4% paraformaldehyde in PBS (pH 7.4, Affimetrix) for 30 minutes at room temperature. The fixation reaction was quenched by the addition of 50 mM NH_4_Cl in DPBS for 10 minutes. <u>For time courses of bacterial infection</u>, bacteria were added at MOI 30 to the cultures for 1 hour at 37°C in 10% CO_2_. After 1 hour of infection, monolayers were washed twice with DMEM without FCS to remove non-binding extracellular bacteria and either harvested directly for the 1-hour time point or incubated with 1 mL of 100 µg/mL gentamycin in DMEM without 10% FCS and incubated for 1 hour to kill non-invaded bacteria. After this incubation, the medium was exchanged for 1 mL DMEM 10% FCS with 15 µg/mL gentamycin until the collection time point. The infected monolayer cultures were collected for qPCR, RNA sequencing, and immunoblot, and the medium was collected for ELISA and Luminex assay at the indicated timepoints after infection.

### Flow cytometry

Following fixation, the cell suspension was incubated with α-MUC1-ED antibody 139H2 (1:100), PE goat α-mouse IgG secondary antibody (1:1,600; 1031-09, SouthernBiotech), and diluted in FACS buffer on ice for 30 minutes in the dark. After antibody incubation, the cell suspension was washed 2 times with ice-cold FACS buffer. Finally, cells were resuspended in ice-cold FACS buffer, and data were collected on a Beckman Coulter CytoFLEX and analyzed with FlowJo software (TriStar).

### Fluorescence microscopy

Following fixation, the HT29-MTX cultures were washed twice with DPBS and permeabilized with DPBS containing 0.2% Triton X-100 for 10 minutes. Non-specific binding sites were blocked with 1% BSA and 22 mg/ml glycine in PBST (DPBS with 0.1% Tween 20) for 30 minutes. Coverslips were incubated with primary antibodies α-MUC1-ED 139H2 or 214D4, α-MUC1-SEA 232A1 (kind gifts from John Hilkens), or α-MUC1-CT antibody CT2 (Thermo, MA5-11202) at 1:100 dilution in PBST with 1% BSA overnight at 4°C or 1.5 hours at RT. After antibody incubation, monolayers were washed 3 times with blocking buffer or DPBS, followed by incubation with secondary antibodies goat α-mouse IgG Alexa Fluor-488/568 (1:100; A11029, A11031; ThermoFisher) or goat α-Armenian hamster IgG Alexa Fluor-488 (1:100; ab173003; Abcam) and DAPI (2 μg/ml; D21490, Invitrogen) diluted in PBST 1% BSA for 1 hour at RT. Coverslips were washed twice with DPBS and twice with MilliQ, dried, and mounted in a droplet of Prolong diamond solution (ThermoFisher) on glass slides. Images were collected on a Leica SPE-II confocal microscope using a 63x objective (NA 1.3, HCX PLANAPO oil) controlled by Leica LAS AF software with default settings to detect DAPI, Alexa488, and Alexa568. Images of Figure 1B were captured on a spinning disk Olympus SpinSR10 system equipped with a Yokogawa W1-SoRa spinning disk mounted on an IX83 stand with an ORCA Flash 4.0 camera (Olympus, Leiderdorp, the Netherlands). The system was run in confocal mode using a 60x oil objective (UPLXAPO XOHR, NA1.5). Z slices imaged at 60x were 0.25 µm to visualize the entire cell layer. Fluorescence was recorded upon sequential excitation by lasers (Coherent, OBIS) with appropriate emission filters and exposure time. The main dichroic mirror was a quadband (D405/488/640nm). Orthogonal views were displayed in CellSens software (Olympus).

### RNA isolation and quantitative reverse transcription PCR

HT29-MTX cultures in 12-well plates infected for 0, 1, 3, 6, or 22 hours were washed once with DPBS and lysed with 1 mL of TRIzol™ (Invitrogen) on ice and stored at −80°C until all time points were collected. RNA was isolated according to the protocol provided by the supplier. In brief, lysed cells were resuspended with 0.2 mL of chloroform per 1 mL of TRIzol™, vortexed thoroughly, and incubated at RT for 2-3 minutes. Next, the solution was centrifuged for 15 minutes at 12,000 g at 4°C, and the colorless upper aqueous phase containing the RNA was carefully transferred to a new tube. 0.5 mL isopropanol per 1 mL of TRIzol™ was added, incubated for 10 minutes at RT, and centrifuged for 10 minutes at 12,000 g at 4°C. Supernatant was discarded, and the RNA pellet was washed once with 1 mL 75% ethanol, centrifuged for 5 minutes at 7500 g at 4°C. The RNA pellet was air-dried for 10 minutes at RT, resuspended in 50 µL RNase-free water, and incubated at 60°C for 10 minutes. RNA concentration was determined using BioDrop, absorbance at 260nm. Two μg of RNA was treated with 2 units DNaseI in reaction buffer with MgCl_2_, in a volume of 18 µl for 30 minutes at 37°C. The reaction was stopped by the addition of 2 µL of 25 mM EDTA and incubation for 10 minutes at 65°C. The RT-qPCR mixture contained 50 ng RNA or MilliQ as a negative control, 0.3 mM final concentration forward and reverse primers, reverse transcriptase (Eurogentec, RT-RTCK-03, 5 U/μL), RNase inhibitor (1 U/μL), 1x No Rox SYBR mastermix blue dTTP (Takyon, UF-NSMT-B0701), completed with DEPC-treated water to a final volume of 20 μl. RT-qPCR primers used were TNFα-Fw 5’-AGCCCATGTTGTAGCAAACCC-3’, TNFα-Rev 5’-GCCTTGGCCCTTGAAGAGGA-3’, IL-8-Fw 5’-CTGGCCGTGGCTCTCTTG-3’, IL-8-Rev 5’-CCTTGGCAAAACTGCACCTT-3’, NONO-Fw 5’-AAAGCAGGCGAAGTTTTCATTC-3’, NONO-Rev 5’-ATTTCCGCTAGGGTTCGTGTT-3’. RT-qPCR reactions were performed using a LightCycler 480 II system (Roche). First, cDNA synthesis was performed at 48°C for 30 minutes, followed by a PCR program with the following steps 1) 95 °C for 5 minutes, 2) 95°C for 5 seconds and 60°C for 30 seconds for 45 cycles and 3) melting curves were generated by incubation at 95°C for 5 seconds, gradually reduced to 65°C in steps of 1 minute, and a final step at 97°C. Results were analyzed using the LightCycler 480 software. Ct values were determined and normalized to the housekeeping gene NONO. Fold changes were calculated using the arithmetic formula (2^−ΔΔCT^).

### IL-8 ELISA assay

Supernatants of infected or TNFα-stimulated cultures were collected in tubes on ice, centrifuged at 12,000 g at 4°C, and stored at −20°C until all the time points were collected. IL-8 concentrations were determined using an IL-8 ELISA kit (R&D Systems, DY208) according to the manufacturer’s instructions. 96-well ELISA plates were coated with 4µg/mL of IL-8 capture antibody overnight at RT. After coating, the plate was washed three times with wash buffer (PBS with 0.05% Tween 20). The plate was then blocked for 1 hour at RT with block buffer (PBS with 1% BSA). The block buffer was removed, and IL-8 standards diluted in reagent diluent, and undiluted samples were added and incubated for 2 hours at RT. Wells were washed twice with washing buffer and incubated with 20 ng/ml detection antibody for 2 hours at RT. Wells were washed twice and incubated with Streptavidin-HRP diluted 1:40 in reagent diluent for 20 minutes at RT. The plate was then washed twice before adding 100 µL substrate solution (ThermoFisher, SB02), followed by incubating for 20 minutes at RT in the dark. 50 µL of the stop solution was added to stop the reaction. Optical density was measured at 450 nm with a FLUOstar Omega microplate reader (BMG Labtech). Sample concentrations in pg/mL were calculated by a 4-parametric curve fit on the standard curve.

### Luminex assay

Supernatants of cultures were collected as described for ELISA. To determine concentrations of 18 selected analytes, multiplex immunoassays were performed at the Multiplex Core Facility (MCF) of the Center for Translational Immunology of the University Medical Center Utrecht, the Netherlands. Immunoassays were developed and validated by the MCF and based on Luminex xMap technology (Luminex, Austin, TX, USA) (58). In brief, samples were incubated with MagPlex microspheres (Luminex) for 1 hour at RT, while continuously shaking, followed by 1 hour of incubation with biotinylated antibodies and 10 minutes of incubation with phycoerythrin-conjugated streptavidin in high-performance ELISA buffer (HPE, Sanquin, Hamburg, Germany). Data acquisition was performed with FLEXMAP 3D equipment in combination with xPONENT software (version 4.3, Luminex), and analyte concentration reported in pg/ml was analyzed by 5-parametric curve fitting using Bio-Plex Manager software (58).

### Western blots

PBS was added to cellular monolayers, and a cell scraper was used to detach the cells. The suspension was transferred to an Eppendorf tube and centrifuged for 5 minutes at 1,500 rpm at 4°C. The pellet was resuspended in DPBS with 1% SDS, and lysis was performed by passing the suspension through a 26G needle approximately 10 times. The suspension was centrifuged at 14,000 g for 5 minutes to remove unlysed material, and the lysate was transferred to a new tube. Protein concentrations were determined using a BCA kit (Pierce). Laemmli sample buffer was added to equal amounts and boiled for 5 min at 95°C. For the detection of MUC1 ED, 30 µg of total protein per sample was run on a 5% mucin gel with wet transfer to a PVDF membrane as previously described (21). For the detection of MUC1-CT, NFκB proteins, and housekeeping proteins, 10 µg of total protein per sample was run on a 10% SDS-PAGE gel. After electrophoresis, semi-dry transfer to PVDF membranes was performed using the Trans-Blot Turbo Transfer system for 7 min at 25 V and 1.3 A. Membranes were blocked with 5% BSA in TSMT (20 mM Tris, 150 mM NaCl, 1 mM CaCl_2_, 2 mM MgCl_2_, adjusted to pH 7 with HCl, with 0.1% Tween 20). Incubation with primary antibodies was performed overnight at 4°C or 1-2 hours at RT with either the α-MUC1-ED antibody 214D4 or 139H2 (1:1000), α-MUC1-CT antibody CT2 (1:1000, Thermo, MA5-11202), or the α-Actin antibody (1:5000; bs-0061R, Bioss), NFκB antibodies (RELA (D14E12, #8242), RELB (D7D7W, #10544), NFKB1 (D4P4D, #13586), NFKB2 (D7A9K, #37359), IκBα (L35A5, #4814), Cell Signaling, 1:1000) diluted in TSMT 1% BSA. Membranes were washed four times with TSMT, followed by incubation with secondary antibodies α-mouse IgG (1:5000, A2304, Sigma) or α-rabbit IgG (1:5000, A4914, Sigma). After antibody incubation, membranes were washed twice with TSMT and twice with TSM for 5 minutes. Blots were developed with the Clarity Western ECL kit (Bio-Rad) and imaged in a Gel-Doc system (Bio-Rad). For the quantification of protein bands, each protein band was first normalized to the corresponding housekeeping protein control then relative expression was calculated either to the corresponding uninfected cells or wild-type cells.

### Library preparation and RNA sequencing

Total RNA was extracted from uninfected and *Salmonella*-infected MUC1-WT, MUC1-ΔCT, and ΔMUC1 cultures as described above. One µg of total RNA per sample was used for mRNA library preparation using Stranded Total RNA Prep, Ligation with Ribo-Zero Plus (Illumina, 20040525) following the manufacturer’s instructions. In brief, samples were subjected to rRNA depletion, and the RNA was first hybridized to DNA probes with Hybridize Probe Master Mix. The mixture was denatured at 95°C for 2 minutes, then cooled to 37°C at a ramp rate of 0.1°C/sec and held for probe hybridization. Following hybridization, rRNA Depletion Master Mix was added to the samples, and the samples were incubated at 37°C for 15 minutes. Probes bound to rRNA were enzymatically digested using Probe Removal Master Mix in a thermal cycler set to 37°C for 15 minutes, followed by 70°C for 15 minutes. RNA was then purified using 60 µL of RNAClean XP beads (Beckman Coulter), washed twice with 80% ethanol, and eluted in elution buffer. Fragmentation and priming were performed by adding Elute, Prime, Fragment 3HC Mix (EPH3) to each sample and incubated at 94°C for 2 minutes. First-strand cDNA synthesis was initiated by adding First Strand Synthesis Master Mix to each fragmented RNA sample. The thermal cycling conditions were: 25°C for 10 minutes, 42°C for 15 minutes, and 70°C for 15 minutes to inactivate the reverse transcriptase. To generate double-stranded cDNA and incorporate dUTP for strand specificity, Second Strand Marking Master Mix (SMM) was added and incubated at 16°C for 1 hour in a thermal cycler and cleaned using AMPure XP beads. DNA was washed twice with 80% ethanol and eluted in Resuspension Buffer. Total double-stranded cDNA was recovered per sample. A-tailing was performed by adding ATL4 (A-Tailing Mix) to each cDNA sample. Samples were incubated at 37°C for 30 minutes, followed by 70°C for 5 minutes. Anchor ligation was achieved by adding RNA Index Anchors (from the IDT for Illumina RNA UD Indexes Set A, Illumina 20040553), Ligation Mix, and Resuspension Buffer to each sample. The mixture was incubated at 30°C for 10 minutes. To terminate ligation, Stop Ligation Buffer was added to each reaction. Adapter-ligated DNA was purified using AMPure XP beads, followed by two washes with 80% ethanol and final elution in Resuspension Buffer. Each cleaned sample was transferred to a new plate for PCR amplification. Amplification of the adapter-ligated cDNA was performed in a PCR reaction containing Enhanced PCR Mix (EPM) and dual-indexed primers (UDP0001– UDP0025) from the Illumina RNA UD Index Set. The thermal cycling protocol was: initial denaturation at 98°C for 30 s, followed by 13–15 cycles of 98°C for 10 s, 60°C for 30 s, and 72°C for 30 s, with a final extension at 72°C for 5 minutes. The amplified libraries were purified with AMPure XP beads and eluted in Resuspension Buffer. Library fragment size distribution and concentration were analyzed using a DNA 1000 chip on the Agilent 2100 Bioanalyzer (Agilent Technologies). Pooled libraries were sequenced by Macrogen Europe (Amsterdam, The Netherlands) on the Illumina NovaSeq platform using 150 bp paired-end reads.

### RNAseq data analysis

RNA sequencing data analysis was performed according to a standardized RNA sequencing analysis pipeline. Briefly, quality control of all sequence reads was checked using FastQC (version 0.11.9) for raw sequence reads and MultiQC (version 1.13 (8dd46e3) for sequences after alignment. Sequencing reads were mapped to the human genome (assembly GRCh38.109) using the STAR RNA aligner (version 2.7.10b). Then, FeatureCounts was used to count the mapped reads (version 2.0.3). For differential expression genes (DEGs) analysis, transcript counts were analyzed using the RStudio (RStudio 2023.03.1+446) package DEseq2 (version 1.46.0) for DEGs, and FDR-corrected values of p < 0.05 were considered statistically significant. PCA data was plotted after batch effect correlation using the limma package (3.48.3) using ggplot2 (3.4.2). Gene set enrichment analysis was done using clusterProfiler (version 4.14.4) with a multiple testing correction method using 0.05 as a significance threshold. Heatmaps were generated using ComplexHeatmaps (version 2.22.0). Volcano plots were generated using EnhancedVolcano (version 1.24.0).

## Data Availability

The RNA sequencing data generated for this study have been deposited in the NCBI Gene Expression Omnibus (GEO) under accession number GSE304522 and are accessible at https://www.ncbi.nlm.nih.gov/geo/query/acc.cgi?acc=GSE304522. Both raw and processed datasets are available.

## Acknowledgements

Microscopy was performed at the Center for Cell Imaging (CCI) of the Faculty of Veterinary Medicine, Utrecht University, and we thank Dr. Richard Wubbolts and Esther van ‘t Veld for their expert advice. We thank the Flow Cytometry and Cell Sorting Facility of the Faculty of Veterinary Medicine at Utrecht University for support. We thank Frank Riemers and Bart Westendorp for their generous help with RNA sequencing analysis. The Luminex assay and analysis were conducted by the MultiPlex Core Facility of the University Medical Center Utrecht, Utrecht, The Netherlands. K. Strijbis has received funding from the European Research Council (ERC) under the European Union’s Horizon 2020 research and innovation program (ERC-2019-STG 852452), by which J. Su was supported.

## Author contributions

Conceptualization: JS, XL, EF, KS. Funding acquisition: KS. Project administration: KS. Methodology: JS, XL, EF, LXZH, KS. Investigation: JS, XL, EF, LXZH. Visualization: JS, KS. Supervision: KS, JPMP. Writing original draft: JS, KS. Review & editing: JS, XL, EF, LXZH, JPMP, KS

---

**Supplementary Figure S1.**
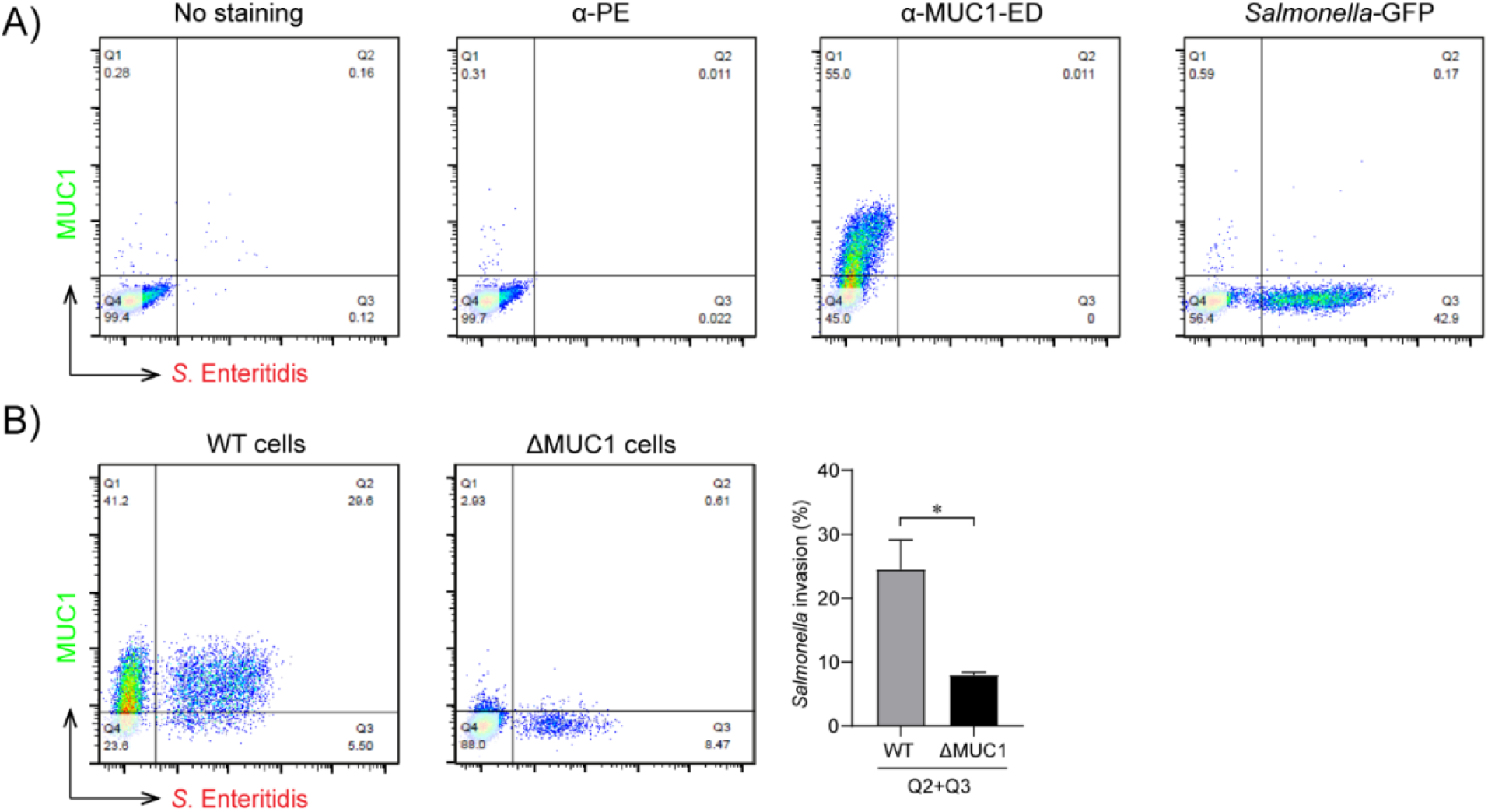
Flow cytometry-based quantification of *Salmonella* infection of intestinal epithelial cells. A) Gating strategy of *Salmonella*-GFP infection of HT29-MTX-WT cells using FACS analysis. Cells were gated based on cells that were not stained, stained with α-PE secondary only, double-stained with α-MUC1-ED antibody 139H2 and α-PE secondary (shown as α-MUC1-ED), and infected with *Salmonella*-GFP only. B) Left, flow cytometry of representative WT and ΔMUC1 cells infected with *Salmonella*-GFP for 1 h and stained with α-MUC1-ED antibody 139H2, and right, total percentage of *Salmonella*-GFP-infected WT and ΔMUC1 cells (Q2+Q3).

## References

1. McCracken VJ, Lorenz RG. 2001. The gastrointestinal ecosystem: A precarious alliance among epithelium, immunity and microbiota. Cellular Microbiology 3:1–11.

2. Linden SK, Sutton P, Karlsson NG, Korolik V, McGuckin MA. 2008. Mucins in the mucosal barrier to infection. Mucosal Immunology 1:183–197.

3. McGuckin MA, Lindén SK, Sutton P, Florin TH. 2011. Mucin dynamics and enteric pathogens. Nature Reviews Microbiology 9:265–278.

4. Hollingsworth MA, Swanson BJ. 2004. Mucins in cancer: Protection and control of the cell surface. Nature Reviews Cancer 4:45–60.

5. Bose M, Mukherjee P. 2019. Microbe – MUC1 Crosstalk in Cancer-Associated Infections 1–13.

6. Van Putten JPM, Strijbis K. 2017. Transmembrane Mucins: Signaling Receptors at the Intersection of Inflammation and Cancer. J Innate Immun 9:281–299.

7. Bramwell ME, Wiseman G, Shotton DM. 1986. Electron-microscopic studies of the ca antigen, epitectin. Journal of Cell Science 86:249–261.

8. Parry S, Silverman HS, McDermott K, Willis A, Hollingsworth MA, Harris A. 2001. Identification of MUC1 Proteolytic Cleavage Sites in Vivo. Biochemical and Biophysical Research Communications 283:715–720.

9. Macao B, Johansson DGA, Hansson GC, Härd T. 2006. Autoproteolysis coupled to protein folding in the SEA domain of the membrane-bound MUC1 mucin. Nat Struct Mol Biol 13:71–76.

10. Hanson RL, Hollingsworth MA. 2016. Functional consequences of differential O-glycosylation of MUC1, MUC4, and MUC16 (Downstream effects on signaling). Biomolecules. MDPI AG 10.3390/biom6030034.

11. Singh PK, Hollingsworth MA. 2006. Cell surface-associated mucins in signal transduction. Trends in Cell Biology 16:467–476.

12. McGuckin MA, Every AL, Skene CD, Linden SK, Chionh YT, Swierczak A, McAuley J, Harbour S, Kaparakis M, Ferrero R, Sutton P. 2007. Muc1 mucin limits both Helicobacter pylori colonization of the murine gastric mucosa and associated gastritis. Gastroenterology 133:1210–1218.

13. Lindén SK, Sheng YH, Every AL, Miles KM, Skoog EC, Florin THJ, Sutton P, McGuckin MA. 2009. MUC1 limits Helicobacter pylori infection both by steric hindrance and by acting as a releasable decoy. PLoS pathogens 5:e1000617.

14. Kato K, Lillehoj EP, Park YS, Umehara T, Hoffman NE, Madesh M, Kim KC. 2012. Membrane-Tethered MUC1 Mucin Is Phosphorylated by Epidermal Growth Factor Receptor in Airway Epithelial Cells and Associates with TLR5 To Inhibit Recruitment of MyD88. The Journal of Immunology 188:2014–2022.

15. Ng GZ, Menheniott TR, Every AL, Stent A, Judd LM, Chionh YT, Dhar P, Komen JC, Giraud AS, Wang TC, McGuckin MA, Sutton P. 2016. The MUC1 mucin protects against Helicobacter pylori pathogenesis in mice by regulation of the NLRP3 inflammasome. Gut 65:1087–1099.

16. Lu W, Hisatsune A, Koga T, Kato K, Kuwahara I, Lillehoj EP, Chen W, Cross AS, Gendler SJ, Gewirtz AT, Kim KC. 2006. Cutting Edge: Enhanced Pulmonary Clearance of Pseudomonas aeruginosa by Muc1 Knockout Mice. The Journal of Immunology 176:3890–3894.

17. Souvannavong V, Saidji N, Chaby R. 2007. Lipopolysaccharide from Salmonella enterica Activates NF-κB through both Classical and Alternative Pathways in Primary B Lymphocytes. Infect Immun 75:4998–5003.

18. Guang W, Ding H, Czinn SJ, Kim KC, Blanchard TG, Lillehoj EP. 2010. Muc1 cell surface mucin attenuates epithelial inflammation in response to a common mucosal pathogen. Journal of Biological Chemistry 285:20547–20557.

19. Lillehoj EP, Hyun SW, Kim BT, Zhang XG, Lee DI, Rowland S, Kim KC. 2001. Muc1 mucins on the cell surface are adhesion sites for Pseudomonas aeruginosa. American Journal of Physiology - Lung Cellular and Molecular Physiology 280.

20. Lillehoj EP, Kim H, Chun EY, Kim KC. 2004. Pseudomonas aeruginosa stimulates phosphorylation of the airway epithelial membrane glycoprotein Muc1 and activates MAP kinase. American Journal of Physiology - Lung Cellular and Molecular Physiology 287.

21. Li X, Bleumink-Pluym NMC, Luijkx YMCA, Wubbolts RW, van Putten JPM, Strijbis K. 2019. MUC1 is a receptor for the Salmonella SiiE adhesin that enables apical invasion into enterocytes. PLoS Pathogens 15:1–20.

22. Gerlach RG, Cláudio N, Rohde M, Jäckel D, Wagner C, Hensel M. 2008. Cooperation of Salmonella pathogenicity islands 1 and 4 is required to breach epithelial barriers. Cellular Microbiology 10.1111/j.1462-5822.2008.01218.x.

23. Chatterjee M, Huang LZX, Mykytyn AZ, Wang C, Lamers MM, Westendorp B, Wubbolts RW, Van Putten JPM, Bosch B-J, Haagmans BL, Strijbis K. 2023. Glycosylated extracellular mucin domains protect against SARS-CoV-2 infection at the respiratory surface. PLoS Pathog 19:e1011571.

24. Chen EY, Tan CM, Kou Y, Duan Q, Wang Z, Meirelles GV, Clark NR, Ma’ayan A. 2013. Enrichr: interactive and collaborative HTML5 gene list enrichment analysis tool. BMC Bioinformatics 14:128.

25. Li X, Wubbolts RW, Bleumink-Pluym NMC, Van Putten JPM, Strijbis K. 2021. The Transmembrane Mucin MUC1 Facilitates β1-Integrin-Mediated Bacterial Invasion. mBio 12:e03491–20.

26. Guang W, Ding H, Czinn SJ, Kim KC, Blanchard TG, Lillehoj EP. 2010. Muc1 Cell Surface Mucin Attenuates Epithelial Inflammation in Response to a Common Mucosal Pathogen. Journal of Biological Chemistry 285:20547–20557.

27. McAuley JL, Linden SK, Png CW, King RM, Pennington HL, Gendler SJ, Florin TH, Hill GR, Korolik V, McGuckin MA. 2007. MUC1 cell surface mucin is a critical element of the mucosal barrier to infection. J Clin Invest 117:2313–2324.

28. Cascio S, Zhang L, Finn OJ. 2011. MUC1 Protein Expression in Tumor Cells Regulates Transcription of Proinflammatory Cytokines by Forming a Complex with Nuclear Factor-κB p65 and Binding to Cytokine Promoters. Journal of Biological Chemistry 286:42248–42256.

29. Ahmad R, Raina D, Joshi MD, Kawano T, Ren J, Kharbanda S, Kufe D. 2009. MUC1-C Oncoprotein Functions as a Direct Activator of the Nuclear Factor-κB p65 Transcription Factor. Cancer Research 69:7013–7021.

30. Bose M, Mukherjee P. 2020. Microbe–MUC1 Crosstalk in Cancer-Associated Infections. Trends in Molecular Medicine 26:324–336.

31. Costa NR, Mendes N, Marcos NT, Reis CA, Caffrey T, Hollingsworth MA, Santos-Silva F. 2008. Relevance of MUC1 mucin variable number of tandem repeats polymorphism in H pylori adhesion to gastric epithelial cells. WJG 14:1411.

32. Carvalho F, Seruca R, David L, Amorim A, Seixas M, Bennett E, Clausen H, Sobrinho-Simoes M. 1997. MUC1 gene polymorphism and gastric cancer–an epidemiological study. Glycoconjugate Journal 14:107–111.

33. Vinall LE, King M, Novelli M, Green CA, Daniels G, Hilkens J, Sarner M, Swallow DM. 2002. Altered expression and allelic association of the hypervariable membrane mucin MUC1 in Helicobacter pylori gastritis. Gastroenterology 123:41–49.

34. Lillehoj EP, Hyun SW, Liu A, Guang W, Verceles AC, Luzina IG, Atamas SP, Kim KC, Goldblum SE. 2015. NEU1 Sialidase Regulates Membrane-tethered Mucin (MUC1) Ectodomain Adhesiveness for Pseudomonas aeruginosa and Decoy Receptor Release. Journal of Biological Chemistry 290:18316–18331.

35. Carson DD. 2008. The Cytoplasmic Tail of MUC1: A Very Busy Place. Sci Signal 1.

36. Raina D, Ahmad R, Kumar S, Ren J, Yoshida K, Kharbanda S, Kufe D. 2006. MUC1 oncoprotein blocks nuclear targeting of c-Abl in the apoptotic response to DNA damage. EMBO J 25:3774–3783.

37. Ren J, Bharti A, Raina D, Chen W, Ahmad R, Kufe D. 2006. MUC1 oncoprotein is targeted to mitochondria by heregulin-induced activation of c-Src and the molecular chaperone HSP90. Oncogene 25:20–31.

38. Radziejewska I, Supruniuk K, Bielawska A. 2021. Anti-cancer effect of combined action of anti-MUC1 and rosmarinic acid in AGS gastric cancer cells. European Journal of Pharmacology 902:174119.

39. Cascio S, Farkas AM, Hughey RP, Finn OJ. 2013. Altered glycosylation of MUC1 influences its association with CIN85: the role of this novel complex in cancer cell invasion and migration. Oncotarget 4:1686–1697.

40. Liu X, Yi C, Wen Y, Radhakrishnan P, Tremayne JR, Dao T, Johnson KR, Hollingsworth MA. 2014. Interactions between MUC1 and p120 Catenin Regulate Dynamic Features of Cell Adhesion, Motility, and Metastasis. Cancer Research 74:1609–1620.

41. Yamamoto M, Bharti A, Li Y, Kufe D. 1997. Interaction of the DF3/MUC1 Breast Carcinoma-associated Antigen and β-Catenin in Cell Adhesion. Journal of Biological Chemistry 272:12492–12494.

42. Wen Y, Caffrey TC, Wheelock MJ, Johnson KR, Hollingsworth MA. 2003. Nuclear Association of the Cytoplasmic Tail of MUC1 and β-Catenin. Journal of Biological Chemistry 278:38029–38039.

43. Li Y, Bharti A, Chen D, Gong J, Kufe D. 1998. Interaction of Glycogen Synthase Kinase 3β with the DF3/MUC1 Carcinoma-Associated Antigen and β-Catenin. Molecular and Cellular Biology 18:7216–7224.

44. Li Y, Kuwahara H, Ren J, Wen G, Kufe D. 2001. The c-Src Tyrosine Kinase Regulates Signaling of the Human DF3/MUC1 Carcinoma-associated Antigen with GSK3β and β-Catenin. Journal of Biological Chemistry 276:6061–6064.

45. Ren J, Li Y, Kufe D. 2002. Protein Kinase C δ Regulates Function of the DF3/MUC1 Carcinoma Antigen in β-Catenin Signaling. Journal of Biological Chemistry 277:17616–17622.

46. Kohlgraf KG, Gawron AJ, Higashi M, Meza JL, Burdick MD, Kitajima S, Kelly DL, Caffrey TC, Hollingsworth MA. 2003. Contribution of the MUC1 tandem repeat and cytoplasmic tail to invasive and metastatic properties of a pancreatic cancer cell line. Cancer Res 63:5011–5020.

47. Ng GZ, Menheniott TR, Every AL, Stent A, Judd LM, Chionh YT, Dhar P, Komen JC, Giraud AS, Wang TC, McGuckin MA, Sutton P. 2016. The MUC1 mucin protects against Helicobacter pylori pathogenesis in mice by regulation of the NLRP3 inflammasome. Gut 65:1087–1099.

48. Mitchell S, Tsui R, Tan ZC, Pack A, Hoffmann A. 2023. The NF-κB multidimer system model: A knowledge base to explore diverse biological contexts. Sci Signal 16:eabo2838.

49. Shih VF-S, Tsui R, Caldwell A, Hoffmann A. 2011. A single NFκB system for both canonical and non-canonical signaling. Cell Res 21:86–102.

50. Banoth B, Chatterjee B, Vijayaragavan B, Prasad M, Roy P, Basak S. 2015. Stimulus-selective crosstalk via the NF-κB signaling system reinforces innate immune response to alleviate gut infection. eLife 4:e05648.

51. Cockram TOJ, Puigdellívol M, Brown GC. 2019. Calreticulin and Galectin-3 Opsonise Bacteria for Phagocytosis by Microglia. Front Immunol 10:2647.

52. Park A-M, Hagiwara S, Hsu DK, Liu F-T, Yoshie O. 2016. Galectin-3 Plays an Important Role in Innate Immunity to Gastric Infection by Helicobacter pylori. Infect Immun 84:1184–1193.

53. Zhao Q, Guo X, Nash GB, Stone PC, Hilkens J, Rhodes JM, Yu L-G. 2009. Circulating Galectin-3 Promotes Metastasis by Modifying MUC1 Localization on Cancer Cell Surface. Cancer Research 69:6799–6806.

54. Piyush T, Chacko AR, Sindrewicz P, Hilkens J, Rhodes JM, Yu L-G. 2017. Interaction of galectin-3 with MUC1 on cell surface promotes EGFR dimerization and activation in human epithelial cancer cells. Cell Death Differ 24:1937–1947.

55. Okamura H, Tsutsui H, Komatsu T, Yutsudo M, Hakura A, Tanimoto T, Torigoe K, Okura T, Nukada Y, Hattori K, Akita K, Namba M, Tanabe F, Konishi K, Fukuda S, Kurimoto M. 1995. Cloning of a new cytokine that induces IFN-γ production by T cells. Nature 378:88–91.

56. Okamura H, Kashiwamura S, Tsutsui H, Yoshimoto T, Nakanishi K. 1998. Regulation of interferon-γ production by IL-12 and IL-18. Current Opinion in Immunology 10:259–264.

57. Bumann D. 2001. Regulated Antigen Expression in Live Recombinant Salmonella enterica Serovar Typhimurium Strongly Affects Colonization Capabilities and Specific CD4^+^-T-Cell Responses. Infect Immun 69:7493–7500.

58. Scholman RC, Giovannone B, Hiddingh S, Meerding JM, Malvar Fernandez B, Van Dijk MEA, Tempelman MJ, Prakken BJ, De Jager W. 2018. Effect of anticoagulants on 162 circulating immune related proteins in healthy subjects. Cytokine 106:114–124.

